# The *Medicago truncatula* nodule-specific cysteine-rich peptides, NCR343 and NCR-new35 are required for the maintenance of rhizobia in nitrogen-fixing nodules

**DOI:** 10.1101/2023.01.23.523609

**Authors:** Beatrix Horváth, Berivan Güngör, Mónika Tóth, Ágota Domonkos, Ferhan Ayaydin, Farheen Saifi, Yuhui Chen, János Barnabás Biró, Mickael Bourge, Zoltán Szabó, Zoltán Tóth, Rujin Chen, Péter Kaló

**Affiliations:** Institute of Plant Biology, Biological Research Centre, Eötvös Lóránd Research Network, Szeged, Hungary; Institute of Genetics and Biotechnology, Hungarian University of Agriculture and Life Sciences, Gödöllő, Hungary; Hungarian Centre of Excellence for Molecular Medicine (HCEMM) Nonprofit Ltd., Szeged, Hungary; Cellular Imaging Laboratory, Biological Research Centre, Eötvös Lóránd Research Network, Szeged, Hungary; School of Life Sciences, Lanzhou University, Lanzhou, China 730000; Cytometry Facility, Imagerie-Gif, Université Paris-Saclay, CEA, CNRS, Institute for Integrative Biology of the Cell (I2BC), 91198, Gif-sur-Yvette, France

**Keywords:** nitrogen-fixing symbiosis, nodule-specific cysteine-rich peptide, symbiosis, indeterminate nodule, *Medicago truncatula*

## Abstract

In the nodules of Inverted Repeat-Lacking Clade legumes, including *M. truncatula*, nitrogen-fixing rhizobia undergo terminal differentiation resulting in elongated and endoreduplicated bacteroids specialised for nitrogen fixation. This irreversible transition of rhizobia is mediated by host produced nodule-specific cysteine-rich (NCR) peptides, of which about 700 are encoded in the *M. truncatula* genome. Some of these NCR peptides, NCR169, NCR211 and NCR247, are essential for nitrogen-fixing symbiosis.

The analysis of bacteroid and symbiotic host cell differentiation revealed that the symbiotic phenotype of *M. truncatula* mutants, *Mtsym19*, *Mtsym20* and NF-FN9363, were defective likewise in the formerly studied *ncr* mutants, *Mtdnf4-1* and *Mtdnf7-2*. The incomplete differentiation of bacteroids triggered premature senescence of rhizobia in the nitrogen fixation zones of mutant nodules.

*Mtsym19* and *Mtsym20* mutants are defective in the same peptide NCR-new35 and the lack of *NCR343* is responsible for the ineffective symbiosis of NF-FN9363.

The activity of *NCR-new35* is significantly lower and limited to the transition zone of the nodule compared with other crucial *NCRs*. The fluorescent protein-tagged version of NCR343 and NCR-new35 localize to the symbiotic compartment. Our discovery added two additional members to the group of *NCR* genes essential for nitrogen–fixing symbiosis in *M. truncatula*.

## Introduction

Legumes can establish nitrogen-fixing endosymbiotic association with soil bacteria, collectively termed rhizobia. This interaction results in the formation of nodules, generally formed on the roots, where rhizobia are hosted and convert atmospheric dinitrogen to ammonia. The interaction is initiated by the mutual recognition of the host legume and rhizobia mediated by signalling molecules of both symbiotic partners (Oldroyd & Downie, 2008). The perception of the bacterial signalling molecules, the nodulation factors (NFs), and the attachment of rhizobia on root hair cells induces two synchronous developmental processes, rhizobial infection and initiation of nodule primordia by activating the mitosis of certain root cortical cells. Trapped bacteria in root hair curls induce the invagination of the plasma membrane forming tubular-like structures with rhizobia inside, termed infection threads (IT). The ITs progress transcellularly towards the nodule primordia and when they reach the dividing root cortical cells, rhizobia are released and colonize many of the nodule cells. Within infected nodule cells, bacteria are encompassed by a plant derived membrane resulting in formation of subcellular compartments called symbiosomes. Bacteria multiply and undergo morphological and metabolic transition to adapt from free-living life-style to their advanced endosymbiotic form termed bacteroids (Jones *et al*., 2007).

Nodules formed on legume roots are classified into two types, the so-called indeterminate or determinate nodules, based on the term of their meristem activity (Hirsch, 1992). The cylindrical shaped indeterminate nodules, such as those found on *Medicago truncatula* roots, possess a persistent meristem during the life-span of the nodules, and as a result, the mature indeterminate nodules consist of a developmental gradient of cells forming distinct zones (Vasse *et al*., 1990). The mitotic activity of meristem cells (zone I, ZI), located at the nodule apex, produces new cells for all nodule tissues. Cells leaving the meristem are relatively small with a large central vacuole (Gavrin *et al*., 2014) and have a four-fold haploid DNA content (4C) (Nagymihaly *et al*., 2017). In these cells in the distal part of the infection zone (ZIId), rod-shaped rhizobia are released from ITs to the host cell cytoplasm and from hereafter, both rhizobia and infected nodule cells undergo differentiation processes through the subsequent nodule zones. The transition of symbiotic cells and rhizobia involves the replication of their genomes without cytokinesis resulting in enlargement of invaded host cells and elongation of bacteria (Vasse *et al*., 1990; Mergaert *et al*., 2006; Nagymihaly *et al*., 2017). The enlarged symbiotic cells in the proximal part of the infection zone (ZIIp) have a DNA content of 8C/16C and their volume is almost occupied by distended vacuoles and symbiosomes containing non-dividing bacteroids. In cells of the first few layers of the nitrogen fixation zone (ZIIId, also termed as transition zone or interzone between ZII and ZIII and referred hereafter IZ in this study), elongated bacteroids surround the collapsed central vacuoles of infected host cells (16C/32C) which are filled with starch granules. Rhizobia and host cells complete their differentiation in IZ and nitrogen fixation begins in the large symbiotic cells having highly endoreduplicated chromosomes (32C/64C) and large volume vacuoles in the nitrogen fixation zone (ZIII). As the indeterminate nodule ages, proximal to the root, a senescence zone (ZIV) evolves wherein the symbiotic interaction terminates and both infected host cells and rhizobia degrade.

Rhizobia, hosted in the nodules of legumes of the Inverted Repeat-Lacking Clade (IRLC) (consisting of legume genera such as *Medicago*, *Pisum*, *Trifolium*, *Vicia*, etc.) and *Aeschynomene* species undergo terminal differentiation (Mergaert *et al*., 2006; Czernic *et al*., 2015). These differentiated bacteroids, reaching 5-10 μm in length and DNA content of 24C in fully differentiated form in *M. truncatula* nodules, are not able to regrow and proliferate outside the indeterminate nodule. The irreversible transition of rhizobia in *M. truncatula* nodules is mediated by host-produced defensin-related nodule-specific cysteine-rich (NCR) peptides (Van De Velde *et al*., 2010). The genome of *M. truncatula* codes for about 700 NCR peptides and similar genes in different numbers have been identified in other IRLC legume species (Mergaert *et al*., 2003; Montiel *et al*., 2017; Zorin *et al*., 2022; Wei *et al*., 2022). Almost all *NCR* genes are specifically expressed in infected cells of *M. truncatula* root nodules and they are activated in successive waves during nodule organogenesis and bacteroid differentiation (Nallu *et al*., 2013; Guefrachi *et al*., 2014). NCR peptides possess highly conserved N-terminal signals which are recognised by the secretion complex of the nodule cells (Wang *et al*., 2010). The secretory pathway delivers the mature NCR peptides, usually 35 to 55 residues in length, to the symbiosomes inducing bacteroid differentiation. The characteristic feature of NCR peptides is the four or six cysteine residues in conserved positions which are presumed to be involved in the formation of intra- and intermolecular disulphide bonds (Mergaert *et al*., 2003). The structural analysis of peptides NCR044 and NCR169 confirmed the presence of different pattern of disulphide bonds that contribute to the structural stability of the peptides (Velivelli *et al*., 2020; Isozumi *et al*., 2021). *M. truncatula* Thioredoxin s1 (Trx s1), which is able to reduce disulphide bonds, is also targeted to symbiosomes and its role has been suggested regulating the activity of NCR peptides by their redox state (Ribeiro *et al*., 2017). These results indicate the importance of cysteine residues and the substitution of the conserved cysteine for serine evidenced the requirement of cysteines and the intracellular disulphide bridges for the symbiotic function of the NCR169 peptide (Horvath *et al*., 2015).

The high number of *NCR* genes in the *M. truncatula* genome implied that they act redundantly. Forward genetic studies of deletion mutants deficient in single NCR peptides, NCR169 or NCR211, revealed that these peptides are essential for bacteroid differentiation and persistence in *M. truncatula* nodules (Horvath *et al*., 2015; Kim *et al*., 2015). Despite these findings, little knowledge is available about the molecular mechanism of NCR peptides mediating bacteroid differentiation *in planta*. A recent reverse genetic analysis of *NCR247* demonstrated that the encoded peptide is required for nodule functioning and provided experimental data about the operation of one of the NCR peptides (Sankari *et al*., 2022).

Biochemical studies demonstrated that NCR247 is able to form a complex with haem facilitating iron uptake by rhizobia. In addition to the prominent positive regulatory role of these NCR peptides in the symbiotic interaction, other peptides, NFS1 and NFS2, control bacterial survival in a strain- and allele-specific manner and negatively regulate the nitrogen-fixing symbiosis in *M. truncatula* (Yang *et al*., 2017; Wang *et al*., 2017).

The *M. truncatula* genome contains several hundreds of *NCR* genes but only few of them have been proven to be required for nitrogen-fixing symbiosis. In this report, we present the phenotypic analysis of three *M. truncatula* symbiotic nitrogen-fixing mutants *Mtsym19*, *Mtsym20* and NF-FN9363. The failure of four different rhizobial strains to colonise the nitrogen fixation zone of the symbiotic nodules indicates that the ineffective symbiotic phenotype is not strain dependent. In the ineffective nodules, the bacteroid differentiation was incomplete suggesting the function of the impaired plant genes in this developmental step. We found that *Mtsym19* and *Mtsym20* are both defective in the same gene, *NCR-new35* and the deletion of *NCR343* in mutant NF-FN9363 resulted in the ineffective symbiotic phenotype.

We show that the substitution of the first cysteine residue either in NCR-new35 or in NCR343 abolishes the symbiotic function of these peptides. Our results indicate that the peptides NCR-new35 and NCR343 are essential for terminal bacteroid differentiation in *M. truncatula* nodules.

## Materials and Methods

### Plant materials, rhizobia strains and growth conditions

Symbiotic mutant NF-FN9363 was identified during symbiotic screens of fast neutron bombarded *M. truncatula* A17 plants (The Samuel Roberts Noble Foundation, Ardmore, Oklahoma, USA). Mutants *Mtsym18* (TR36), *Mtsym19* (TR183) and *Mtsym20* (TRV43) were obtained following gamma ray irradiation of *M. truncatula* cv. Jemalong (J5) (Sagan *et al*., 1995; Morandi *et al*., 2005). *M. truncatula* cv. Jemalong or genotype A17 were used as a wild-type (wt) controls for characterization of the symbiotic phenotype of *Mtsym19*, *Mtsym20* and FN-NF9363 mutants. Symbiotic mutants were crossed to *M. truncatula* A20 for genetic mapping. Seeds were germinated as described in the *M. truncatula* handbook (Garcia *et al*., 2006) Seedlings were grown under symbiotic conditions in a zeolite substrate (Geoproduct Kft., Mád, Hungary) in growth chambers and kept under a photoperiod of 16 h of light / 8 h of darkness at 24 °C. Four-day-old pre-grown plants were inoculated with the rhizobium strains *Sinorhizobium* (*Ensifer*) *medicae* WSM419, ABS7 or *S. meliloti* 1021 carrying the pXLGD4 plasmid expressing the *hemA∷lacZ* reporter gene or strain *S. meliloti* FSM-MA carrying the pMEpTrpGUSGFP plasmid expressing the β-glucuronidase-GFP fusion protein. Rhizobia were cultured in tryptone-yeast (TY) media (pH 7.2) supplemented with 6 mM CaCl_2_ (Beringer, 1974). The bacterial liquid culture was pelleted by centrifugation and resuspended in Gibson liquid media (Gibson & Nutman 1960) at a final dilution 1:50 (~OD_600_ 0.1) and used for inoculation. Plants were harvested or scored for symbiotic phenotype indicated in each experiment.

### Histological analyses and microscopy

The rhizobium-dependent nodulation phenotype of the ineffective mutant and wild-type plants was analysed at 3 weeks post inoculation (wpi) with different bacterial strains using a Leica MZ10 F stereo microscope at 2.5x standard magnification.

### *lacZ* and SYTO13 staining

To assess bacterial colonization, mutant and wt plants were inoculated with rhizobia strains carrying the pXLGD4 plasmid and 14-day-old nodules post inoculation were harvested in 1xPBS buffer (pH 7.4), vacuum-fixed in 4% (w/v) paraformaldehyde solution for 3 x 30 sec in an Eppendorf 5301 concentrator and post-fixed on ice for 30 min. Fixed nodules were embedded in 5% (w/v) agarose and 65 μm thick longitudinal sections were prepared using a Leica VT1200S vibrotome. In order to visualize the rhizobium-infected cells, nodule sections were stained for β-galactosidase activity in X-Gal staining solution (Bovin *et al*., 1990) for 30 minutes at 37 °C. Stained sections were analysed using an Olympus BX41M light microscope (Olympus Life Science Europa GmbH, Hamburg, Germany) with 10x objective and images were taken with an Olympus E-10 digital camera. The symbiotic nodule cell structure and bacteroid morphology were also analysed on nodule sections stained with 5 μM SYTO13 (Thermo Fisher Scientific, diluted in 1xPBS buffer pH 7.4) for 20 min at room temperature and imaged using a Leica TCS SP8 confocal laser scanning microscope. The microscope configuration was as follows: objective lenses HCX PL FLUOTAR 10x dry (NA: 0.3) and HC PL APO CS2 63x oil (NA: 1.4); zooms: 0.75 and 2.14; excitation: OPSL 488 nm laser (SYTO13, green) and OPSL 552 nm laser (red autofluorescence); spectral emission detector: 510-540 nm (SYTO13) and 558 nm – 800 nm (red autofluorescence).

### GUS reporter assay

The promoter activity of the *NCR* genes was assayed with the promoter-*GUS* reporter gene constructs generated using the Gateway cloning technology (Invitrogen). The cloned promoter fragments (819 bp of *pr NCR-new35*, 1620 bp of *prNCR343*, 1178 bp of *prNCR169* and 2055 bp of *prNCR211*) of the four *NCR* genes were recombined from pDONR201 entry vector constructs into a modified Gateway destination vector pKGWFS7 containing the dsRED fluorescent transformation marker under the control of a constitutively active *Arabidopsis thaliana* ubiquitin10 (pUBQ10) promoter (pKGWFS7-pUBQ10∷dsRED). The constructs were introduced into the roots of *M. truncatula* A17 plants using the *Agrobacterium rhizogenes*-mediated hairy root transformation carried out as described by Boisson-Dernier and co-workers (2001). To detect GUS activity, 65 μm thick fixed sections of nodules, harvested at 2 and 3 wpi with *S. medicae* WSM419, were infiltrated with X-Gluc staining solution (1 mM 5-bromo-4-chloro-3-indolyl-β-D-glucuronic acid substrate, 50 mM sodium phosphate buffer (pH 7.4), 2.5 mM potassium ferrocyanide, 2.5 mM potassium ferricyanide, 20% methanol and Triton X-100 at 0.1 ml/100 ml) for 30-40 min at 37 °C. GUS stained nodule sections were analysed and imaged similar to *lacZ*-stained nodule sections.

### Scanning electron microscopy

Nodule samples were treated and sections were prepared for scanning electron microscopy as described earlier (Domonkos *et al*., 2017).

### Genetic mapping and allelism test

The F2 segregation populations were generated by self-pollination of the F1 progeny of the crosses of the NF-FN9363, *Mtsym19* and *Mtsym20* mutant lines, respectively with the *M. truncatula* A20 genotype. To determine the map position of the mutant loci, linkage analyses were carried out with the genotypes of the F2 individuals determined for a marker set of *M. truncatula* (Choi *et al*., 2004; Mun *et al*., 2006). Primers of additional markers developed for fine mapping and to identify deletion borders are listed in Table S1. To carry out allelism test between *Mtsym19* and *Mtsym20*, F3 plants homozygous for both symbiotic loci and carrying the opposite parental homozygous alleles for the PCR-based MtB267 marker were selected and crossed. The success of the cross was confirmed by genotyping of the offspring and the allelic relationship was assayed by analysing their nodulation phenotype and growth habit. The genomic DNA of *M. truncatula* plants was isolated using the ZenoGene40 plant DNA purification kit (Zenon Bio, Szeged, Hungary).

### Identification of deletions in the genome of *M. truncatula* symbiotic mutants

The database of copy number variations identified with whole genome array-based comparative genomic hybridization (CGH) analysis of *M. truncatula* fast neutron bombarded mutant lines (https://medicago-mutant.dasnr.okstate.edu/mutant/index.php) were mined to identify mutagenesis events (In/Del and SNP variations) co-segregating with the symbiotic mutant locus of FN-NF9363.

To detect genetic alterations in the genome of symbiotic mutants *Mtsym18* (TR36), *Mtsym19* (TR183) and *Mtsym20* (TRV43), an analysis of RNAseq data was carried out. Nodules of the symbiotic mutants were harvested at 2 wpi with *S. medicae* WSM419 (pXLGD4) in liquid nitrogen, total RNA was extracted with TRI Reagent (Sigma) and purified with Direct-zol RNA MiniPrep Kit (Zymo Research, Irvine, CA, USA). RNA samples were treated with DNaseI on Zymo-Spin columns (Zymo Research) according to the manufacturer’s instructions to remove the genomic DNA. For RNA sequencing, the quality of total RNA was tested on an Agilent 2100 bioanalyzer followed by ribosomal RNA depletion using the Ribominus plant kit (Invitrogen). Libraries were prepared from 1.2 μg rRNA-depleted RNA using the Illumina TruSeq RNA Sample Preparation Kit v2 and sequenced using Illumina NextSeq 500/550 with a minimum of 25 million reads in 150 bp paired-end mode. RNA sequence analysis was carried out using the CLC Genomics workbench (9.5.3) software. Raw reads (36 million for TRV43 and 35 million for TR183) from the Illumina sequencing were trimmed (~5% of the raw reads were discarded) to remove the adaptors and low quality ambiguous nucleotides (Quality limit 0.05, ambiguous limit 2). Trimmed reads were mapped to mRNA reference sequence subsets obtained from the *M. truncatula* A17r5.0 genome assembly (MtrunA17r5.0-ANR-EGN-r1.6.cds) https://medicago.toulouse.inra.fr/MtrunA17r5.0-ANR/). The number of reads of the symbiotic mutants were analysed between genetic markers Chr4g0020111 and Chr4g0020631.

### Protein sequence analysis

Multiple alignments of NCR peptide sequences were created using the CLC Genomics Workbench 9.5.3 program with default settings. The phylogenetic analysis was conducted on the Phylogeny.fr online platform (http://www.phylogeny.fr/simple_phylogeny.cgi). The alignment of the sequences was performed with the MUSCLE program (v3.8.31), the phylogenetic tree was constructed using the maximum likelihood method using the PhylML program (v3.1/3.0 aLRT) and visualized by TreeDyn (v198.3).

### Gene expression analysis

The transcriptional activity of *NCR-new35, NCR211, NCR343, NCR169* as well as senescence- and defence-related genes was analysed with reverse transcription quantitative polymerase chain reaction (RT-qPCR). The expression of *NCR* genes was monitored in wt nodules harvested at 2 and 3 wpi with *S. medicae* WSM419. For the expression analysis of senescence- and defence-related genes, nodules of wt and *nad1-3*, NF-FN9363, *Mtsym20*, *Mtdnf4-1* and *Mtdnf7-1* mutants were harvested at 3 wpi with *S. medicae* WSM419. RNA was extracted in the same way as for RNAseq analysis. Total RNA was quantified on Nanodrop-1000 spectrophotometer and cDNA was prepared from 1 μg total RNA with SuperScript III Reverse Transcriptase (Life Technologies, Invitrogen) using oligo-dT primers according to the manufacturer’s protocol. RT-qPCR reactions were performed on a LightCycler 96 (Roche, Mannheim, Germany) using qPCRBIO SyGreen Mix (PCR Biosystems, London, UK) according to the manufacturer’s instructions. Relative expression of the target genes was normalized using the housekeeping gene *Polypyrimidine tract-binding protein 2* (*PTB*, MtrunA17_ Chr3g0126461) and a gene (MtrunA17_Chr3g0126781) with an ubiquitin domain as reference genes. No RT reaction and the intron sequence of gene MtrunA17_Chr3g0126781 was used to test cDNA samples for genomic DNA contamination. Data were analysed with the LightCycler 96 SW1.1 software. Primer sequences used for RT-qPCR are listed in Table S1.

### Analysis of elongation and DNA content of rhizobia

To measure the bacteroid elongation, rhizobia were isolated from nodules at 16 days post inoculation (dpi). Nodules collected in 1x PBS (pH 7.4) buffer were crushed and homogenized with a sterile glass pestle, the debris was pelleted with centrifugation for 10 min at 2000 rpm at 4 °C and the upper phase was filtered through 50 μm and 20 μm filters (Sysmex/Partec CellTrics filter). Filtered bacteria were pelleted by centrifugation at 7000 rpm for 15 min at 4 °C and resuspended in 20 μl 1xPBS buffer. Purified bacteroids were stained in 20 μM propidium iodide (PI) solution diluted in 1xPBS buffer (pH 7.4) for 30 min and the images were captured at 40x magnification using Leica TCS SP8 confocal laser scanning microscopy. The microscope configuration was as follows: objective lens PL Fluotar 40x oil (NA: 1); zoom: 0.75; excitation: OPSL 552 nm laser; spectral emission detector: 573 - 696 nm. The length of bacteroids was measured on the microscope images using the ImageJ software.

The DNA content and the size of the bacteroid populations were analysed by flow cytometry. Nodules were chopped in BEB buffer (125 mM KCl, 50 mM Na-succinate in 50 mM N-[Tris(hydroxymethyl)methyl]-2-aminoethanesulfonicacid sodium salt (TES) buffer (pH 7), 1% BSA) for bacteroid extraction, and filtered through a 50-μm nylon mesh. Bacteroids or free-living bacteria suspensions were heat-shocked at 70°C for 30 min, and labelled for 10 min with 50 μg/ml of PI, a DNA fluorochrome intercalating dye. Data acquisition was made using a Cytoflex S cytometer (Beckman Coulter). DNA content of 50,000 – 100,000 stained bacteria/bacteroids was measured by 561 nm excitation, through a 610/20 nm band-pass filter. Population was first gated on an SSC-Area/PI-Area dot plot; doublets were then discarded on a PI-Area/PI-Height dot plot. Finally, DNA content or Forward Scatter (FSC) was measured on their respective histograms.

### Complementation experiments using hairy-root transformation

Constructs for genetic complementation experiments were generated using either single-site or multisite Gateway Recombination Cloning Technology (Thermo Fisher Scientific). The PCR amplified products containing the native promoter fragments, *NCR* gene and native 3’ UTR sequences (*prNCR343-1620bp∷NCR343gDNA-3’UTR-543bp, prNCR345-1705bp∷NCR345gDNA-3’UTR-732bp, prNCR341-1659bp∷NCR431gDNA-3’UTR-853bp* and *prNCR-new35-819bp∷NCR-new35gDNA-3’UTR-902bp*) were cloned into pKGW-RR binary destination vector from entry clones generated in pDONR201 vector. The *NCR344* gene construct was generated in pKm43GW-rolD∷EGFP destination vector by recombining the 1476 bp long promoter region, the coding sequence and 472 bp long 3’UTR fragment cloned into entry vectors pDONRP4-P1R, pDONR221 and pDONRP2R-P3, respectively. To investigate the effect of the substitution of the first cysteine residues to serine in the mature peptide of NCR343, NCR-new35 and NCR211, constructs were generated in pKm43GW-rolD∷EGFP destination vector. The 1620, 819 and 2055 bp long promoter regions of *NCR343, NCR-new35* and *NCR211* were inserted into vector pDONRP4P1, respectively. The fragments coding for NCR343-C34S, NCR-new35-C41S and NCR211-C28S were generated by overlapping extension PCR. The *NCR211* coding sequence containing the Cys-Ser substitution was cloned into pDONR221 vector including the first and second exons with the translational Stop codon and a part of the subsequent second intron (*NCR211-mod.coding 1-2.exon-STOP-2.intron - 449bp*). The amplified fragment of the rest of the second intron and the third exon together with the 3’UTR were cloned into pDONRP2R-P3 vector (*NCR211 2.intron 1081bp-3.exon-3’UTR 618bp*). The modified coding sequence with the native 3’UTR of *NCR343* was generated in pDONR221 vector (*NCR343-mod.coding-3’UTR 543bp*) and *NCR-new35* (*NCR-new35-mod.coding-3’UTR 289bp*): The third entry clones in pDONRP2R-P3 vectors carried additional 3’UTR sequences (*NCR343-3’UTR 252bp; NCR-new35-3’UTR 613bp*), respectively. Fragments for the entry clones were amplified with Phusion DNA polymerase (ThermoFisher) or CloneAmpTM HiFi PCR Premix (Takara Bio) using the corresponding primers listed in TableS1. Constructs were introduced into the mutant and wt plants using *A. rhizogenes*-mediated hairy root transformation (Boisson-Dernier *et al*., 2001). The transgenic roots were selected based on the GFP or dsRED fluorescence using a Leica MZ10F fluorescence stereomicroscope.

### Subcellular localization of NCR peptides

To investigate the localization of NCR343 and NCR-new35 peptides in symbiotic nodule cells, constructs coding for mCherry tagged versions of the peptides were generated in pK7m34GW-rolD∷EGFP destination vector using Multisite Gateway Technology (Thermo Fisher Scientific). The entry clones contained the same promoter fragments in pDONRP4-P1R which were used for complementation experiments. The coding sequences of the NCR343 and NCR-new35 peptides from the translational start site without stop codon were inserted into pDONR221 and the mCherry fluorescent tag was cloned into the vector pDONRP2R-P3. The fragments of the entry vectors were recombined into the destination vector pK7m34GW-rolD∷EGFP. The constructs were introduced into NF-FN9363 (pNCR343∷NCR343-mCherry) and *Mtsym20* (pNCR-new35∷NCR-new35-mCherry) mutant plants with *A. rhizogenes*-mediated hairy-root transformation. The signal of fluorescence-tagged NCR343 and NCR-new35 peptides was analysed at 4 wpi on non-fixed 65 μm thick longitudinal and 5 μM SYTO13 stained nodule sections using a Visitron spinning disk confocal microscope equipped with a Yokogawa CSU-W1 spinning disk unit, an Andor Zyla 4.2 Plus camera and an Olympus IX83 inverted microscope (Visitron Ssystem GmBH, Puchheim, Germany). For imaging SYTO13, 488 nm laser excitation and 500-550 nm emission range was used. The mCherry signal was imaged with 561 nm laser excitation and 570-640 nm emission detection range. For low magnification nodule imaging 10x UPlanFL N dry objective (NA: 0.30) and for closeups 100x UPLXAPOXO (NA: 1.45) oil objective was used. Composite images were created with ImageJ program.

## Results

### Nodules of *Medicago truncatula* deletion mutants are defective in colonization of the nitrogen fixation zone by rhizobia

The NF-FN9363, *Mtsym19* (TR183) and *Mtsym20* (TRV43) ineffective symbiotic (Fix-) mutants were identified formerly in genetic screens for symbiotic nitrogen fixation mutants of fast neutron bombarded (NF-FN9363) or γ-ray irradiated (*Mtsym19* and *Mtsym20*) *M. truncatula* A17 (NF-FN9363) or Jemalong (*Mtsym19* and *Mtsym20*) populations (Xi *et al*., 2013; Sagan *et al*., 1995; Morandi *et al*., 2005). *Mtsym19* and *Mtsym20* were reported representing distinct symbiotic loci (Morandi *et al*., 2005) but based on the results described below, we found that these mutants are allelic and therefore only one allele was used in some experiments. The NF-FN9363, *Mtsym19* and *Mtsym20* ineffective mutants were inoculated with different rhizobia strains during the mutant screens. To test the strain dependence of the phenotype of these ineffective symbiotic mutants, the nodulation and growth phenotype of wild-type and NF-FN9363, *Mtsym20, Mtdnf4-1* (*dnf4-1*) and *Mtdnf7-2* (*dnf7-2*) (Starker *et al*., 2006; Domonkos *et al*., 2013) mutant plants were analysed 3 wpi with *S. meliloti* strains 1021 and FSM-MA and *S. medicae* strains ABS7 and WSM419. Mutant plants showed the symptoms of nitrogen starvation (yellow leaves, reduced growth) 3 wpi with all the tested rhizobia strains, even with the highly compatible *S. medicae* WSM419 strain (Terpolilli *et al*., 2008) (Figure 1a). Mutant plants developed roundish or slightly cylindrical white nodules, indicating the absence of leghemoglobin, with each tested rhizobia strain (Figures 1b and S1). Nitrogen deficiency of the ineffective symbiotic mutants induced the development of increased number of nodules and also resulted in reduced dry weight of the aerial part of mutant plants compared with wild-type plants at 3 wpi with each rhizobia strain (Figure S1b). To analyse the colonization of symbiotic nodules, mutant and wild-type plants were inoculated with *S. medicae* WSM419 (pXLGD4), which constitutively expresses the *lacZ* gene (*hemA∷lacZ*). Longitudinal sections of nodules were stained for β-galactosidase activity 14 dpi and the presence of rhizobia was analysed by light microscopy. Wild-type nodules showed the characteristic zonation of indeterminate nodules and were colonized with rhizobia in zones IId, IIp, II-III (IZ) and III at this time point (Fig. 1b lower panel). Mutant nodules were reduced in size compared with wild-type ones and in addition, the zones of indeterminate nodules could be distinguished in mutant nodules. Rhizobia-infected nodule cells were observed in the infection zone (ZIId and ZIIp) and interzone of mutant nodules indicating that the function of the impaired symbiotic genes is essential for the differentiation or the persistence of differentiated rhizobia in mutants NF-FN9363, *Mtsym19* and *Mtsym20* (Fig. 1b lower panel). The nodule colonization in mutants NF-FN9363, *Mtsym19* and *Mtsym20* was similar to that in *M. truncatula dnf4* and *dnf7* mutants defective in NCR peptides NCR211 and NCR169, respectively (Kim *et al*., 2015; Horvath *et al*., 2015). Therefore, and for the reasons described below, the mutants NF-FN9363, *Mtsym19* and *Mtsym20* were analysed along with the symbiotic mutants *Mtdnf4-1* and *Mtdnf7-2*.

**Fig. 1.**
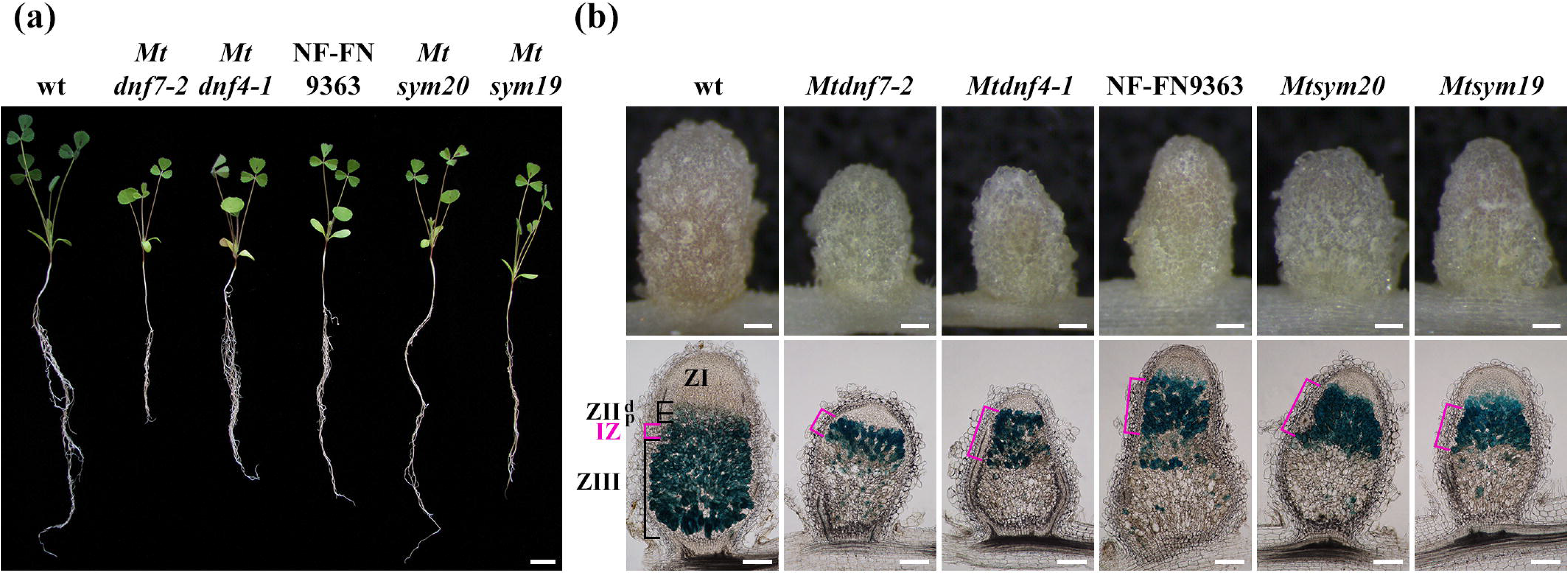
The ineffective nodulation phenotype of *M. truncatula* symbiotic mutants *Mtdnf7-2*, *Mtdnf4-1*, NF-FN9363, *Mtsym20* and *Mtsym19*. (a) Ineffective nitrogen-fixing (Fix-) symbiotic mutants showed the symptoms of the nitrogen starvation (retarded growth, pale green leaves) compared with wild-type (wt) plant two weeks following inoculation with strain *Sinorhizobium medicae* WSM419 carrying the *lacZ* reporter gene. (b) Slightly elongated white nodules were formed on the roots of the Fix-symbiotic mutant plants and cylindrical pink nodules were observed on wt roots 14 days post inoculation (dpi) with *S. medicae* WSM419 (upper images). Longitudinal nodule sections stained for β-galactosidase activity show the normal colonization of nodule zones characteristic for wt indeterminate nodules but in ZIII of Fix-nodules are devoid of rhizobia infected cells (lower images). The zones were labelled with square brackets and the IZ was highlighted in magenta. ZIId, distal part of infection zone; ZIIp, proximal part of infection zone; IZ, interzone; ZIII, nitrogen fixation zone; Bars, (a) 2 cm; (b) 200 μm.

To analyse nodule colonization in more detail, mutant and wild-type nodule sections were stained with nucleic acid-binding dye SYTO13 at 14 dpi with *S. medicae* WSM419 and examined by laser-scanning confocal microscopy that enabled the visualization of stained bacteria and plant nuclei fluoresced in green, the autofluorescent cell wall and accumulated polyphenolic compounds (red channel) simultaneously (Figure 2). Nodules of wild-type plants showed rhizobia-colonized cells in the infection zone (ZIId, ZIIp), interzone (IZ) and nitrogen fixation zone (ZIII) (Figure 2a1-a8). The bacterial invasion of ZII and IZ of mutant and wild-type nodules was similar, albeit the interzone, where bacteria complete their differentiation into symbiotic form, was more extensive and enlarged in mutant nodules compared with wild-type nodules (Figure 1b and Figure 2a1-f1). In contrast to wild-type nodules, the cells in nitrogen fixation zones of mutant nodules were not colonized by rhizobia but showed sporadic autofluorescence (Figure 2b1-f1 and b8-f8). Autofluorescence in nodule tissues often arises from phenolic-compounds derived from pathogen-induced lignification that generally results in robust and extensive defence responses which were characteristics of some *M. truncatula* symbiotic mutants (Bourcy *et al*., 2013; Wang *et al*., 2016; Domonkos *et al*., 2017). In contrast, the early induced decomposing process, the premature senescence is frequently observed in ineffective symbiotic nodules (Van de Velde *et al*., 2006). Early senescence generates less intensive and sporadic autofluorescence that was found previously in *Mtdnf4-1* and *Mtdnf7-2* nodules (Kim *et al*., 2015; Horváth *et al*., 2015). In addition to the difference in the extension and intensity between the defence-reactions and premature senescence-related autofluorescence, the two responses can be distinguished based on activation of marker genes specific for each process. The transcriptional activities of senescence and defence response-specific marker genes were monitored in NF-FN9363, *Mtsym20*, *Mtdnf4*, *Mtdnf7-2*, the *nad1-3* (*nodules with activated defense 1-3;* Domonkos *et al*.,) mutant and Jemalong wild-type nodules at 3 wpi using RT-qPCR. A strong induction of cysteine proteinase genes *MtCP2* and *MtCP6* associated with nodule senescence in *M. truncatula* (Guerra *et al*., 2010) was detected in the ineffective symbiotic nodules of all studied mutants (Figure S2a). In contrast, marker genes of defence responses, a chitinase (*MtCHI*) and *MtPR10* (Class-10 *PATHOGENESIS-RELATED PROTEIN*) genes were not induced in the mutants NF-FN9363, *Mtsym20*, *Mtdnf4* and *Mtdnf7-2* but showed high transcriptional activity in the *M. truncatula nad1-3* mutant (Figure S2b and c), that controls defence-like responses in symbiotic nodules, indicating that the autofluorescence was the result of premature senescence in the nitrogen fixation zones of NF-FN9363, *Mtsym19, Mtsym20*, *Mtdnf4-1* and *Mtdnf7-2* nodules (Figure 2b8-f8).

**Fig. 2.**
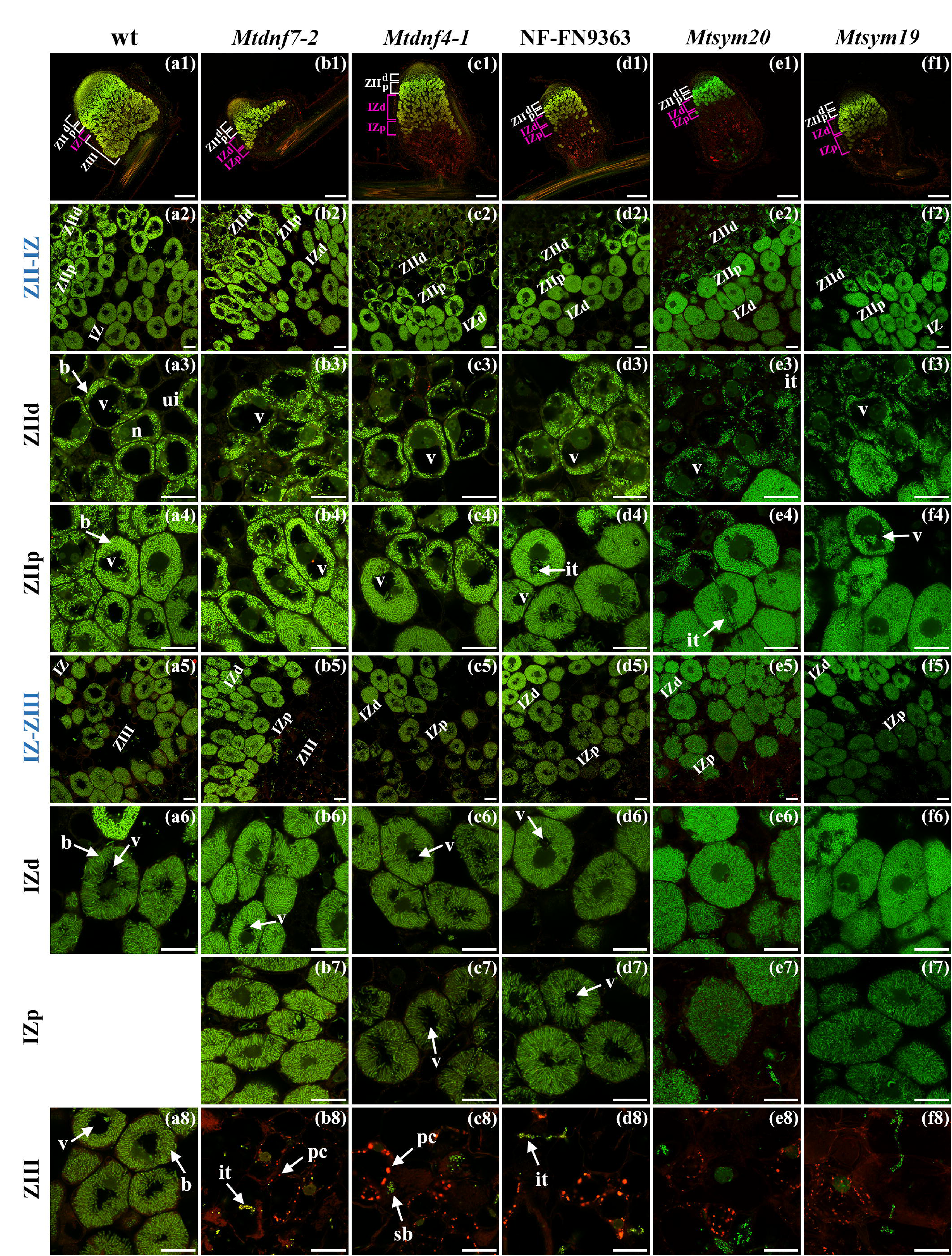
Laser confocal microscopic analyses of symbiotic cell structure and bacteroid morphology on SYTO13 stained nodule sections of the Fix-mutants NF-FN9363, *Mtsym19* and *Mtsym20* compared with nodules of wt and *Mtdnf7-2*, *Mtdnf4-1* plants 14 days post inoculation with *S. medicae* WSM419. (a1-f1) SYTO13-stained longitudinal nodule sections show the defect of bacterial invasion in ZIII of ineffective symbiotic nodules. The green channel visualizes SYTO13-stained bacteroids and plant cell nuclei, autofluorescence is pseudocolored in red. Higher magnification shows the transition between ZII and IZ (a2-f2) and IZ-ZIII (a5-f5). (a3-f3; a4-f4) No obvious alteration was observed in bacterial occupation of infected cells or the morphology of bacteroids in distal part of ZII (ZIId) between wt and mutant nodule cells. (a6-f6; b7-f7) Moderate elongation of bacteroids was found in the proximal part of IZ (IZp) cells of mutant nodules compared with wt nodule cells. This region is composed of a single cell layer in wt nodules but it was extended in mutant nodules. Mutant infected cells in IZp contain disordered bacteroids indicating the inception of their disintegration. (a8-f8) The cells in ZIII but its first cell layer (IZd) in mutant nodules (b7-f7) were devoid of bacteroids and contained autofluoresced speckles indicating the presence of early senescence-related phenolic compounds. Bars, (a1-f1) 200 μm; (a2-f8) 20 μm. it, infection thread; b, bacteroid; sb, saprophytic bacteria; v, vacuole; n, nucleus; ui, uninfected cells.

Higher magnification of SYTO13-stained nodule sections revealed that infected cells in ZII and IZ exhibited similar morphology in mutant and wild-type nodules (Figure 2a1-f5). Bacterial release was normal in the distal part of ZII (ZIId) of mutant nodule cells (Figure 2 a3, b3, c3, d3, e3 and f3) and rhizobia commenced to differentiate in the proximal part of ZII (ZIIp) (Figure 2a4, b4, c4, d4, e4 and f4). These nodule cells in ZIIp, characterized by moderately elongated rhizobia with intense fluorescence and enlarged vacuoles, were clearly visible in wild-type (Figure 2a2 and a4), *Mtdnf7-1* (Figure 2b2 and b4) and *Mtdnf4-1* (Figure 2c2 and c4) nodules but vacuole extension was less pronounced in nodules of FN9363 (Figure 2d2 and d4), *Mtsym20* (Figure 2e2 and e4) and *Mtsym19* (Figure 2f2 and f4). Cells in the first layer of the nitrogen fixation zone showed similar morphology, possessing compressed vacuoles, in wild-type and all mutant nodules although rhizobia were slightly shorter and disorganized in mutant symbiotic cells (Figure 2a5, a6, b5, b6, c5, c6, d5, d6, e5, e6, f5 and f6). Although this region is composed of a single layer of cells in wild-type nodules (Gavrin *et al*., 2014; Figure 2 a1), we observed the extension of this part of nodules in all mutants (Figure 2b1, c1, d1, e1, and f1) indicating the arrest of further differentiation of infected nodule cells. Symbiotic cells with large vacuoles and differentiated bacteroids oriented towards these vacuoles were observed in the nitrogen fixation zone of wild-type nodules (Figure 2a6 and a8). In contrast, vacuoles remained reduced in IZ cells of mutant nodules, resembling to the morphology of cells with compressed vacuoles in interzones (Figure 2b7, c7, d7, e7 and f7). Mutant nodules did not contain infected cells in the region corresponding to the mature nitrogen fixation zone of wild-type nodules but sporadic autofluorescence and non-differentiated rod shape saprophytic bacteria released from the infection threads were detected occasionally in this zone (Figure 2a8, b8, c8, d8, e8 and f8).

### Bacteroid elongation and morphology are impaired in NF-FN9363 and *Mtsym20* nodules

Rhizobia undergo morphological changes including cell enlargement and genome amplification in the indeterminate nodules of *M. truncatula* (Mergaert *et al*., 2006) resulting in elongated (E-type) bacteroids (Montiel *et al*., 2017). The morphology of rhizobia in the interzone cells of mutant nodules indicated the impaired elongation of bacteroids (Figure 2b5-7, c5-7, d5-7, e5-7 and f5-7). In order to determine the level of differentiation, the length and ploidy level of bacteria were measured. Bacteria were isolated from mutant and wild-type nodules 14 dpi, stained with PI and imaged by confocal laser scanning microscopy. The size of at least 1200 bacterial cells was measured using the ImageJ software and the relative ratio of bacteria in different size ranges was compared between wild-type and mutant nodules. Bacterial populations isolated from mutant nodules showed a shift to the smaller size range compared with wild-type sample indicating an imperfect elongation of bacteroids in *Mtdnf4-1*, *Mtdnf7-2*, NF-FN9363 and *Mtsym20* nodules (Figure 3a).

**Fig. 3.**
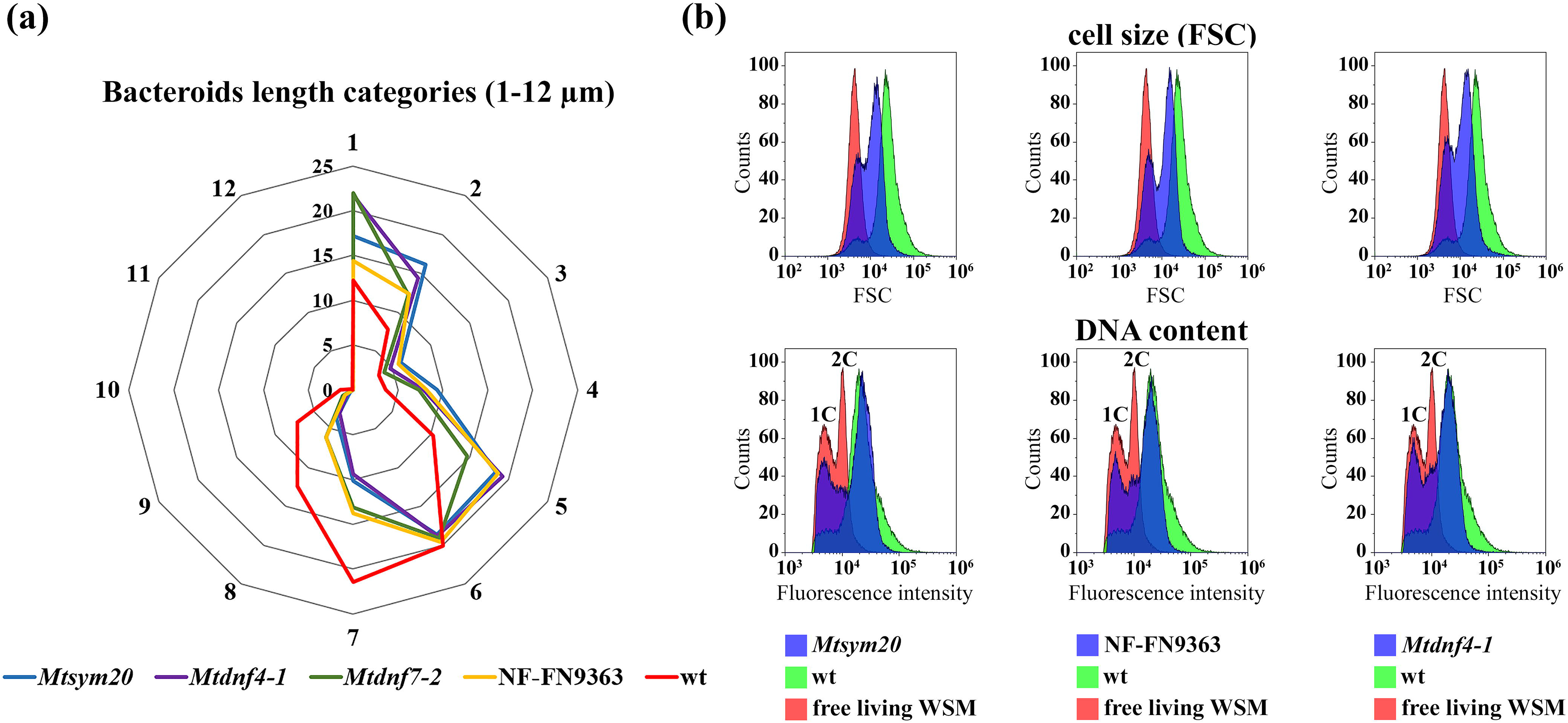
Bacteroid differentiation is similarly impaired in NF-FN9363 and *Mtsym20* mutant nodules as in the formerly identified ineffective symbiotic mutants *Mtdnf4-1* and *Mtdnf7-2* at 2 weeks post inoculation with *S. medicae* WSM419. (a) The distribution of size of bacteroids isolated from wt and mutant nodules at 2 wpi, stained with PI, captured by confocal laser scanning microscopy and measured on the images using the ImageJ program. The size of bacteroids from mutant nodules had a higher proportion of shorter bacterial cells compared with wild-types and rhizobia above 6 μm in length were hardly observed in mutant nodules although these classes in wt nodules were evident. The size of at least 1200 bacterial cells was measured and their relative distribution in length classes is presented. Values around the radar chart indicate bacterial length (μm) and circles show percentage (%). (b) The relative size and DNA content of *S. medicae* WSM419 bacteroids isolated from wt and mutant nodules at 16 dpi measured by flow-cytometry. Bacteroid relative size was determined by the forward light scatter (FSC) and endoreduplication was measured by fluorescence intensity. The size and DNA content of *S. medicae* WSM419 bacteroid populations isolated from mutant nodules shifted to smaller size and lower ploidy level compared with bacteroids found in wt nodules.

The size and DNA content of rhizobia isolated from *Mtdnf4-1*, NF-FN9363 and *Mtsym20* mutant and wild-type nodules were further analysed by flow cytometry. Isolated bacteroids and cultured bacteria were purified and stained with PI. The analysis of FSC intensity, that is proportional to the size of bacteroids, showed that the majority of bacteroids isolated from wild-type nodules were enlarged compared with cultured rhizobia (Figure 3b). The bacterial population purified from mutant nodules contained a mixture of smaller sized cells peaked at the free-living rhizobia and at smaller sized bacteriods compared with wild-type samples. This finding was consistent with length measurements of bacteriods which showed the higher proportion of shorter rhizobia in mutant nodule cells compared with bacteria isolated form wild-type nodules (Figure 3a). The mean DNA content of bacteroids isolated from mutant nodules was higher than the peak of 1C or 2C ploidy levels of cultured *S. medicae* WSM419 cells indicating that bacteroids isolated from mutant nodules were polyploids (Figure 3b). The peaks of the DNA content of rhizobia purified from mutant and wild-type nodules were very similar suggesting an advanced genome amplification of bacteroids in mutant nodules. However, the population of bacterial cells isolated from mutant nodules was narrower at a higher fluorescent range compared with rhizobia purified from wild-type nodules indicating fewer numbers of bacteroids with amplified genomes at the highest degree. The analysis of bacterial size and DNA content of rhizobia showed the similar positive correlation between genome amplification and enlargement of bacteroids detected earlier (Mergaert *et al*., 2006). To detect alterations in mutant nodules in more detail, the ultrastructure of wild-type and NF-FN9363, *Mtsym20*, *Mtdnf4-1* and *Mtdnf7-2* mutant nodules was analysed by scanning electron microscopy (SEM) 18 dpi with *S. medicae* WSM419. Consistent with the light and fluorescent microcopy images, no visible difference was detected in the morphology and invasion of nodule cells in the distal part of ZII of wild-type and mutant nodules (Figure S3a2, b2, c2, d2, e2, a3, b3, c3, d3 and e3). However, rhizobia around the vacuoles did not fill up completely the infected cells in the proximal part of IZ of mutant nodules and this discontinuity between the rhizobia and plant cell walls probably indicates the induction of degradation of cytoplasm and rhizobia (Figure S3a4, b4, c4, d4 and e4). The shrinking of cell content and the aggregation of rhizobia was more apparent in the first cell layer in IZ of mutant nodules compared with wild-type nodule cells, which were fully packed with rhizobia, indicating the advanced collapse of infected mutant cells (Figure S3b5-e5). Close-up images of free-living (Figure S3f1) and symbiotic rhizobia in ZIId and ZIIp of wild-type nodules showed rough-surfaced rhizobia (Figure S3a2 and a3) that probably corresponds to the folding of peribacteroid membranes observed in this region on transmission electron microscopy images of bacteroids of a former study (Vasse *et al*., 1990). As bacteroid differentiation progresses, this morphology changes and smooth-surfaced rhizobia are detected in the IZ and ZIII in wild-type nodules (Figure S3a4 and a6). This alteration of bacterial surface was apparent in the distal part of IZ of all mutant nodules (IZd; Figure S3b4, c4, d4 and e4) implying that bacteroids differentiate similarly in IZ cells of mutant and wild-type nodules. However, disorder and aggregation of bacteroids, often tangled with cell debris, were observed in the proximal IZ cells of mutant nodules (IZp; Figure S3b5, c5, d5 and e5) compared with healthy infected cells of wild-type nodules containing elongated bacteroids orientated towards the vacuoles (Figure S3a4). This disintegration, characterized by vacuolated cells and degraded bacteroids, was more advanced in the few infected cells of ZIII of mutant nodules indicating that decayed cell content was resorbed (Figure S3b6, c6, d6 and e6). These data reveal that rhizobia are able to differentiate to bacteroids in *Mtdnf4-1*, *Mtdnf7-2*, NF-FN9363 and *Mtsym20* nodules but their complete transition is either disrupted or the persistence of bacteroids is defective in mutant nodules.

### Ineffective symbiotic *Medicago truncatula* mutants NF-FN9363, *Mtsym19* and *Mtsym20* are defective in *NCR* genes

Genetic mapping combined with exploiting the high-density genome array-based comparative genomic hybridization (aCGH) platform of *M. truncatula* (Chen *et al*., 2017) or facilitated with transcriptome analysis of symbiotic mutants were applied to identify the impaired genes in symbiotic mutants NF-FN9363, *Mtsym19* and *Mtsym20*. Mutant plants were crossed to *M. truncatula* genotype A20 and genetic mapping was carried out using F2 segregating populations.

The mutant locus of NF-FN9363 was positioned on chromosome 6 (LG 6) below the genetic marker Crs towards MtB178 (Figure S4a). To facilitate the identification of gene defective in NF-FN9363, the genomic positions of copy number variations detected in the mutant line using the array-based CGH (https://medicago-mutant.dasnr.okstate.edu/mutant/index.php) were analysed. A pile of probesets corresponding to the genomic position of the symbiotic locus of NF-FN9363 indicated a potential large deletion in the mutant genome. The deletion was verified by PCR-based markers that defined an almost 500 kb deletion in the genome of NF-FN9363 (Figure 4A and S5, Table S1).

**Fig. 4.**
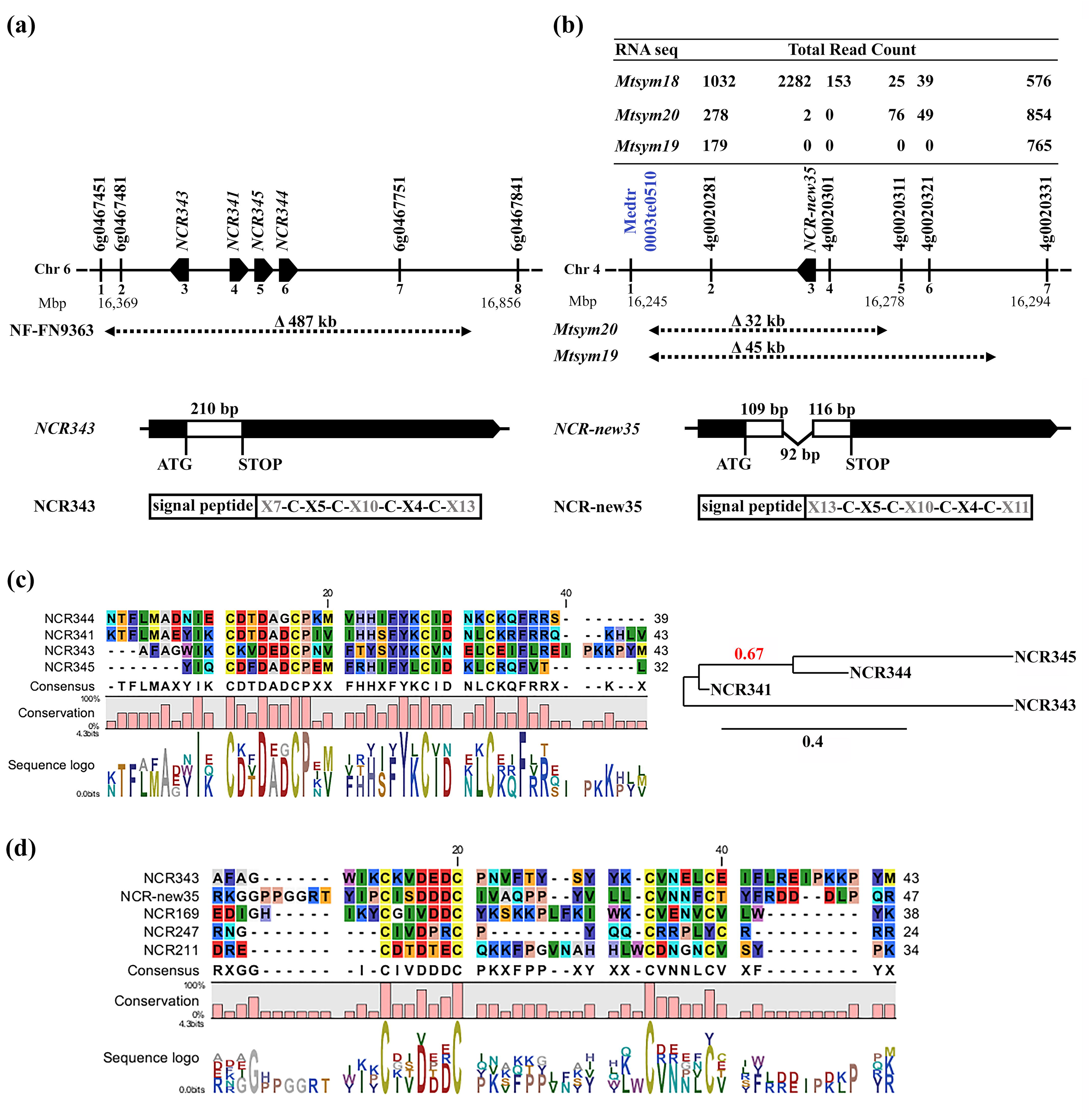
The identified deleted regions and the gene models in the symbiotic loci of NF-FN9363 and *Mtsym19*, *Mtsym20* mutants. (a) Genetic mapping and high-density genome array-based comparative genomic hybridization (aCGH) identified a nearly 500 kb deletion including four *NCR* genes in the NF-FN9363 genome. (b) The extension of the deleted regions in the mutant loci of *Mtsym19* and *Mtsym20* was determined by PCR based markers and by analysing reads obtained from RNAseq. The structure of the *NCR343* (a) and *NCR-new35* (b) genes (coding sequences, the intron of *NCR-new35* and 5’ UTRs and 3’ UTRs) are represented by white boxes, thin excised black line and thick black lines, respectively) and the features of the encoded NCR343 (a) and NCR-new35 (b) peptides containing a signal peptide and mature peptides with four cysteine residues in conserved positions. (c) Multiple sequence alignment and phylogenetic analysis of mature peptides encoded by the *NCR343*, *NCR341, NCR345* and *NCR344* genes deleted in the symbiotic locus of NF-FN9363 and (d) the sequence analysis of mature NCR peptides proved to be crucial for differentiation and persistence of rhizobia during nitrogen-fixing symbiosis. The degree of sequence conservation of the residues between the four mature peptides is represented below the alignment. Colours are based on Robert Fletterick’s “Shapely models.”

The mutant loci of *Mtsym19* and *Mtsym20* were positioned in the same genomic region on chromosome 4 (LG 4) between genetic markers 4g0020111, 4g0020421 and 4g0020631 (Figure S4a). Former genetic analysis suggested that *Mtsym19* and *Mtsym20* belong to different complementation groups (Morandi *et al*., 2005). However, the similar position of the mutant loci of *Mtsym19* and *Mtsym20* indicated either the defect of two different neighbouring genes or the malfunction of the same gene in the two mutants. In order to verify their allelic relationship, an allelism test was carried out using F3 mutant plants selected from the populations of *Mtsym19* x A20 and *Mtsym20* x A20 crosses. Two F3 maternal homozygote plants for the PCR-based genetic marker MtB267 were selected from the *Mtsym19* mapping population and two F3 plants showing paternal homozygous genotype for the same genetic marker were chosen from the *Mtsym20* mapping population and crossed to generate hybrid plants. The different genotypes of the parental lines allowed us to test heterozygosity of the progeny of the F3 mutant plants. The progeny of the three successful crossings were all Fix– (n=8) indicating that *Mtsym19* and *Mtsym20* belong to the same complementation group (Figure S4b).

We presumed that the mutations in *Mtsym19* and *Mtsym20* affected the level of transcripts of the genes impaired in the mutant plants. In order to facilitate gene identification, the transcript abundance in nodules of *Mtsym19, Mtsym20* and the symbiotic mutant *Mtsym18*, belonging to an independent symbiotic complementation group (Morandi *et al*., 2005), was analysed at 2 wpi with *S. medicae* WSM419. Filtered sequence reads were mapped against the *M. truncatula* genome assembly (A17r5.0) and the number of reads was analysed in the genomic region between genetic markers 4g0020111 and 4g0020631. Absent or reduced number of reads aligned to a short non-specific sequence, were detected between gene models 4g0020281 and 4g0020321 in *Mtsym19*, and between 4g0020281 and 4g0020301in *Mtsym20* compared with *Mtsym18*, indicating deletions in these genomic regions (Figure 4B and S6a). Further analysis by PCR amplifications identified a less than 45-kb and a less than 32 kb deletion in the *Mtsym19* and *Mtsym20* genomes, respectively. The deletion in *Mtsym20* spanned three genes encoding for two putative proteins and a Nodule-specific Cysteine-Rich (NCR) peptide (Figure S6b).

### Deletion of gene *NCR343* is responsible for the symbiotic defect of mutant NF-FN9363

The deletion in NF-FN9363 removed more than 30 predicted genes or gene models including four *NCR* genes, *NCR341* (6g0467551), *NCR343* (6g0467511), *NCR344* (6g0467581) and *NCR345* (6g0467571) which were the primary candidates responsible for the symbiotic phenotype of NF-FN9363 (Figure S5b). All four *NCR* genes code for peptides with four cysteines in conserved positions and the sequence analysis of the four mature NCR peptides revealed that NCR341, NCR344 and NCR345 showed higher similarity to each other compared with NCR343 (Figure 4c). The most similar mature peptides of NCR341 and NCR344 are both cationic (pI =8.67 and 7.95), the mature peptide NCR345 is anionic (pI = 4.59), while the cleaved product of NCR343 results in a slightly anionic (pI = 6.33) mature peptide. In order to define that the lack of what gene or genes caused the symbiotic phenotype of NF-FN9363, genetic complementation experiments were carried out. The genomic fragments of the genes *NCR341*, *NCR343*, *NCR344* and *NCR345* containing at least 1.5 kb native promoters and native 3’ UTRs were introduced into NF-FN9363 roots using *A. rhizogenes*-mediated hairy root transformation. Transformed roots were detected by either DsRed or green fluorescent protein markers (Figure S7). The roots were inoculated with *S. medicae* WSM419 (pXLGD4) and nodules were stained for β-galactosidase activity at 4 wpi. Nodule cells in nitrogen fixation zone of mutant NF-FN9363 were devoid of rhizobia (Figure 1b), therefore we assessed the complementation of nodules based on the presence of rhizobia in ZIII. White undeveloped nodules with colonized cells by rhizobia in ZII and IZ were detected on roots transformed with empty vector or with the genes *NCR341*, *NCR344* and *NCR345* indicating that these *NCR* genes were not able to restore the effective symbiotic interaction in NF-FN9363 (Figure 5a and Figure S7g1-i3 and l1-m). The nodules developed on the roots of NF-FN9363 transformed with the construct of *NCR343* were elongated and pink, and the bacterial invasion of ZIII in these nodules was similar to wild-type nodules, suggesting that these were functional nodules on NF-FN9363 roots (Figure 5a and Figure S7f1-f3 and k1-k3). The rescue of the symbiotic phenotype of NF-FN9363 with gene *NCR343* indicated that the loss of this gene caused the ineffective symbiotic phenotype of mutant NF-FN9363.

**Fig. 5.**
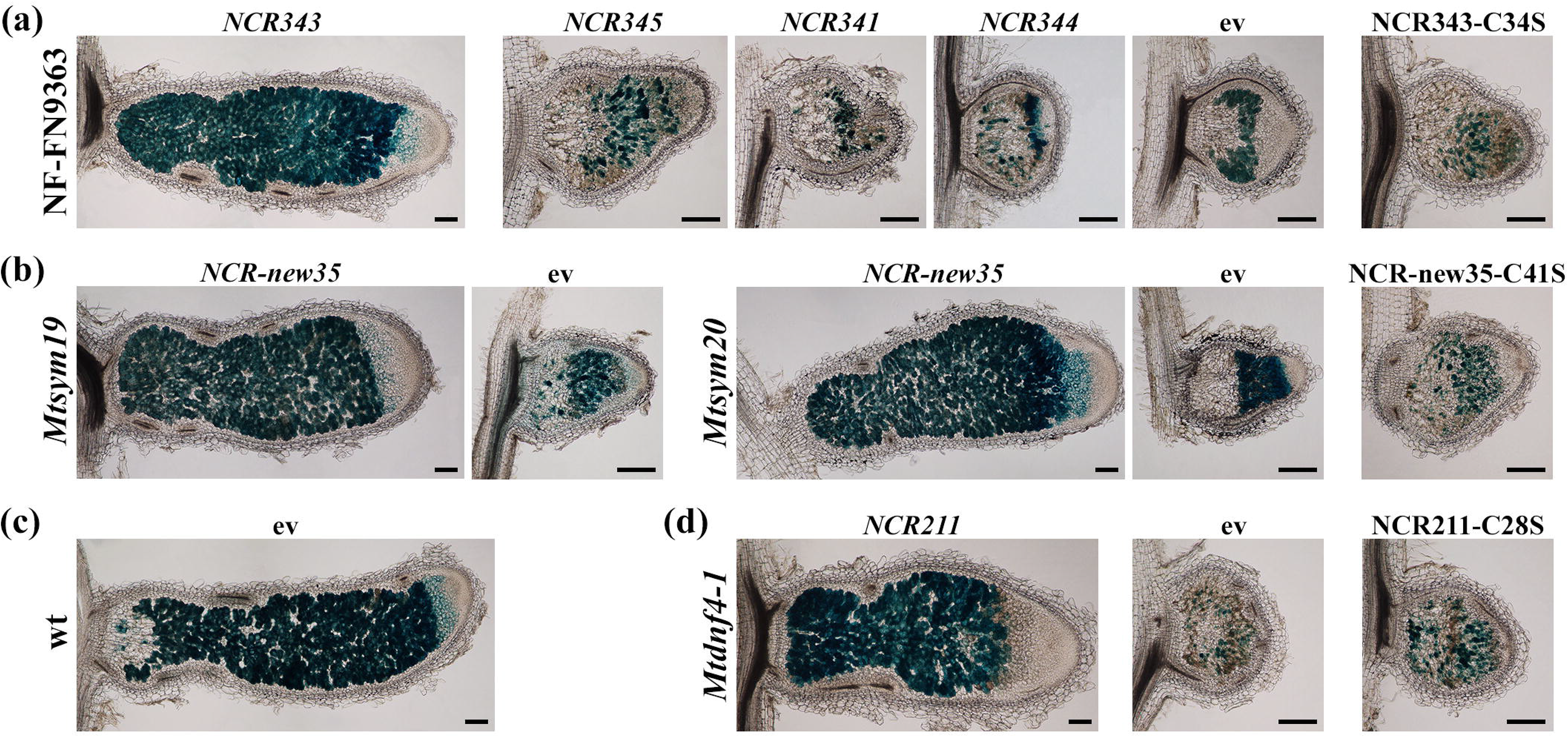
*NCR343* and *NCR-new35* are required for development of functional nodules and the first cysteine residues of NCR343 and NCR-new35 peptides are essential for their function. The rescue of the nodulation defect of mutants NF-FN9363, *Mtsym19* and *Mtsym20* was identified based on the restoration of morphology and colonization of nodules formed on hairy roots transformed with empty vector or *NCR* peptide genes and inoculated with *S. medicae* WSM419 carrying the *lacZ* marker gene. Longitudinal sections of 4-week-old nodules were stained for β-galactosidase activity. The nodules on NF-FN9363 roots transformed with *NCR343* (a), on *Mtsym19* and *Mtsym20* roots transformed with *NCR-new35* (b), on wt roots transformed with an empty vector (c) and on *Mtdnf4-1* roots transformed with *NCR211* genes (d) showed characteristic zonation of indeterminate nodules indicating the restoration of the symbiotic phenotypes. Gene constructs of *NCR345*, *NCR341*, and *NCR344* did not restore the symbiotic phenotype of nodules developed on transformed roots of NF-FN9363 indicating that these peptides are not essential for the symbiotic interaction between *M. truncatula* A17 and *S. medicae* WSM419. The substitution of the first cysteine residue for serine abolished complementation in the mutant NCR343 (a), NCR-new35 (b) and NCR211 (d) genes indicating the essential role of the first cysteines. Bars, (a–d), 200 μm.

### The lack of *NCR-new35* is responsible for the ineffective symbiotic phenotype of *Mtsym19* and *Mtsym20*

The identified deletions in mutants *Mtsym19* and *Mtsym20* overlapped by about 32 kb removing three genes including the gene *NCR-new35*, formerly termed *NCR014* (Bang *et al*., 2017) (Figure 4b and S6B), which made *NCR-new35* the best candidate for *MtSYM19* and *MtSYM20*. In order to confirm that the deletion of *NCR-new35* caused the ineffective symbiotic phenotype of *Mtsym19* and *Mtsym20*, roots of the two mutants were transformed with the wild-type *NCR-new35* gene controlled by an 819 bp native promoter fragment and inoculated with *S. medicae* WSM419 (pXLGD4). DsRed-fluorescent transgenic *Mtsym19* and *Mtsym20* roots expressing the gene *NCR-new35* developed elongated and pink nodules (Figure S7b1-b3, d1-d3 and o1-o3) in contrast to white round-shaped or slightly cylindrical nodules formed on empty vector-transformed roots of mutant plants (Figure S7c1-c3, e1-e3 and p1-p3). In order to visualize bacterial invasion, longitudinal sections of 4-week-old nodules were stained for β-galactosidase activity. Rhizobia-colonized cells were only observed in zones II and IZs of nodules formed on empty vector-transformed *Mtsym19* and *Mtsym20* roots (Figure 5b). Nodules developed on mutant roots transformed with the *NCR-new35* construct showed the typical zonation of indeterminate nodules with colonized cells in the nitrogen fixation zone like nodules on empty vector-transformed wild-type roots (Figure 5b, 5c and Figure S7a1-a3, j1-j3). The restoration of nodule colonization in ZIII of mutant nodules confirmed that gene *NCR-new35* corresponds to *MtSYM19* and *MtSYM20*.

### Replacement of a single cysteine residue in NCR211, NCR343 and NCR-new35 abolishes their symbiotic function

NCR peptides usually contain four or six cysteine residues in conserved positions (Alunni *et al*., 2007). We previously demonstrated that the replacement of a single or multiple cysteine residues of the symbiotic peptide NCR169 ceased its activity *in planta* indicating that each cysteine residue is essential for the symbiotic function of NCR169 (Horvath *et al*., 2015). To confirm the requirement of cysteine residues for the function of NCR-new35, NCR343 and NCR211, we introduced constructs coding for modified peptides, wherein the first cysteines were substituted for serines, into the roots of *Mtsym20*, NF-FN9363 and *Mtdnf4* mutant plants, respectively using *A. rhizogenes*-mediated hairy root transformation. The nodules on the roots of mutants transformed with the modified *NCRs* driven by their native promoters were small and white, suggesting that these were malfunctioning nodules (Figure 5). These nodules did not show the typical zonation of indeterminate nodules and the proximal regions of the nodules corresponding to the nitrogen fixation zones were devoid of bacteria, indicating that the first cysteine residues are essential for the function of NCR-new35, NCR343 and NCR211.

### *NCR-new35* is expressed low in symbiotic cells compared with *NCR169*, *NCR211* and *NCR343*

The members of the large family of *NCR* genes in *M. truncatula* are almost exclusively expressed in symbiotic nodule cells and they are activated in successive waves during nodule differentiation (Guefrachi *et al*., 2014). In order to investigate the expression of genes *NCR-new35* and *NCR343*, their activity was monitored with the *GUS* reporter gene and RT-qPCR analysis and we also analysed the nodule transcriptome data of different *M. truncatula* nodule zones obtained by laser-capture microdissection (LCM) (Roux *et al*., 2014).

To analyse and compare the expression pattern of *NCR-new35* and *NCR343* with the previously identified *NCR169* and *NCR211* genes essential for effective nitrogen fixation (Horvath *et al*., 2015; Kim *et al*., 2015), the promoters of these *NCR* genes were fused to the β-glucuronidase (*GUS*) reporter gene and the constructs were introduced into wild-type *M. truncatula* roots using *A. rhizogenes*-mediated hairy root transformation. Nodules were subjected to histochemical staining for GUS activity at 2 and 3 wpi with *S. medicae* strain WSM419. *NCR343*, *NCR211* and *NCR169* promoters showed activity predominantly in the interzone of two-week-old nodules (Figure 6c-e). In contrast, a low level of *GUS* expression was restricted mainly to the first few cell layers of the interzone and even weaker activity was found sporadically in the distal part of nitrogen fixation zone when the reporter gene was driven by the promoter of *NCR-new35* (Figure 6b). The activity of these promoters is in agreement with LCM data (Roux *et al*., 2014), confirming that *NCR-new35*, *NCR343*, *NCR211* and *NCR169* are mainly active in the IZs of two-week-old nodules and *NCR-new35* is expressed at a much lower level compared with the other three *NCR* genes (Figure 6). The analysis of the promoter activities with the *GUS* reporter gene at 3 wpi revealed that the expression of *NCR169* and *NCR343* and to a less extent the activity of *NCR211* extended to the nitrogen fixation zone (Figure 6g-j). In order to further validate their expression in nodules, we monitored the transcript levels of *NCR-new35*, *NCR343*, *NCR211* and *NCR169* with RT-qPCR in nodules harvested at 2 and 3 wpi. In agreement with the LCM transcriptome data, *NCR343*, *NCR211* and *NCR169* genes showed high relative expression compared with *NCR-new35* (Figure 6k). These transcriptome and *GUS* activity data indicate that *NCR-new35* shows different spatial activity and lower expression intensity compared with the other three essential NCR peptide genes.

**Fig. 6.**
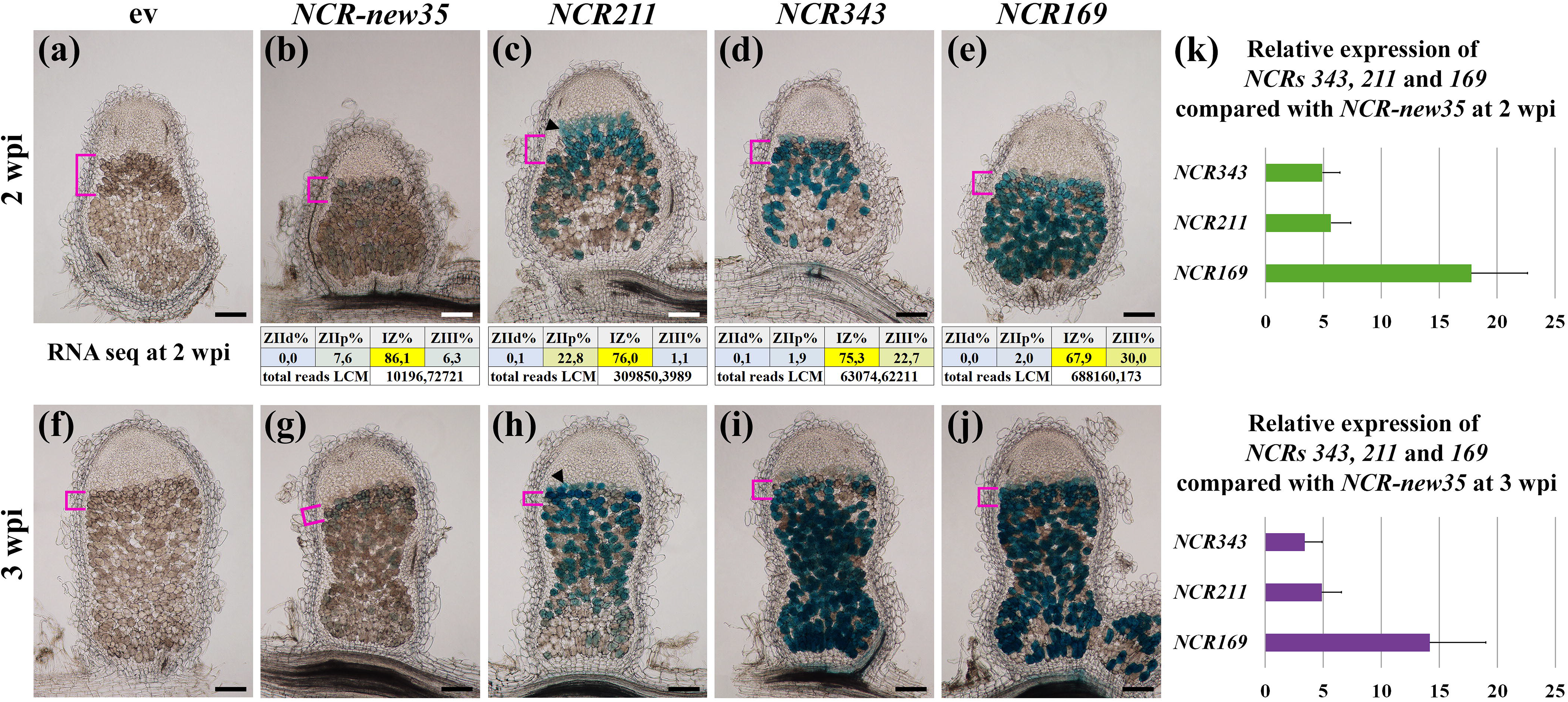
The expression analysis of the *NCR-new35* and *NCR343* genes is compared with the activity of *NCR211* and *NCR169*. The empty vector (a and f) and the construct of promoter fragments of *NCR-new35* (b and g), *NCR211* (c and h), *NCR343* (d and i) and *NCR169* (e and j) genes fused to the β-glucoronidase (GUS) gene, respectively were introduced into the roots of wt Jemalong plants with *A. rhizogenes*-mediated hairy-root transformation. Nodule sections were stained for GUS activity at two (a-e) and three (f-j) wpi with *S. medicae* WSM419. The relative spatial expression of *NCR-new35*, *NCR211*, *NCR343* and *NCR169* genes 2 wpi with *S. medicae* WSM419 generated with RNA sequencing of different nodule zones obtained with laser-capture microdissection (LCM) are shown below the images of 2-week-old nodules. (k) Relative expression of *NCR169*, *NCR211* and *NCR343* compared with *NCR-new35* was analysed in wild-type nodules by reverse transcription quantitative polymerase chain reaction (RT-qPCR) 2 and 3 wpi with rhizobia. Magenta square brackets, IZ; black arrowhead, ZIIp. Bars, (a-j) 200 μm.

### NCR-new35 and NCR343 peptides localize to symbiosomes

NCR peptides usually have a conserved signal peptide which cleavage is essential for targeting of NCR peptides to the bacteroids (Wang *et al*., 2010). Previous studies demonstrated the localization of NCR169 and NCR211 peptides to the bacteroids in the symbiotic cells in the IZ and nitrogen fixation zone (Horvath *et al*., 2015; Kim *et al*., 2015). To make certain the subcellular localization of NCR-new35 and NCR343 peptides, translational fusions to the *mCherry* reporter gene driven by native *NCR-new35* or *NCR343* promoters were generated and introduced into *Mtsym20* and NF-FN9363 mutant roots, respectively using hairy-root transformation. The functional complementation of the mutant nodules indicated that the fusion proteins retained their activity and the fluorescent tag did not perturb the function of NCR-new35-mCherry and NCR343-mCherry proteins (Figure 7). The red fluorescence of NCR-new35-mCherry and NCR343-mCherry proteins was detected in the IZ and nitrogen fixation zone. The signal of the NCR-new35-mCherry fusion protein is partially contradictory to the expression pattern of *NCR-new35* detected at 3 wpi with rhizobia (Figure 6g) and points to the sustained stability of the NCR-new35-mCherry fusion protein in the nitrogen fixation zone. In contrast to NCR-new35-mCherry, the subcellular localization of NCR343-mCherry corresponded completely to the spatial expression pattern of *NCR343* (Figure 6i). Higher magnification of symbiotic cells showed co-localization of mCherry-tagged NCR-new35 and NCR343 with SYTO13-stained rhizobia (Figure 7c and f).

**Fig. 7.**
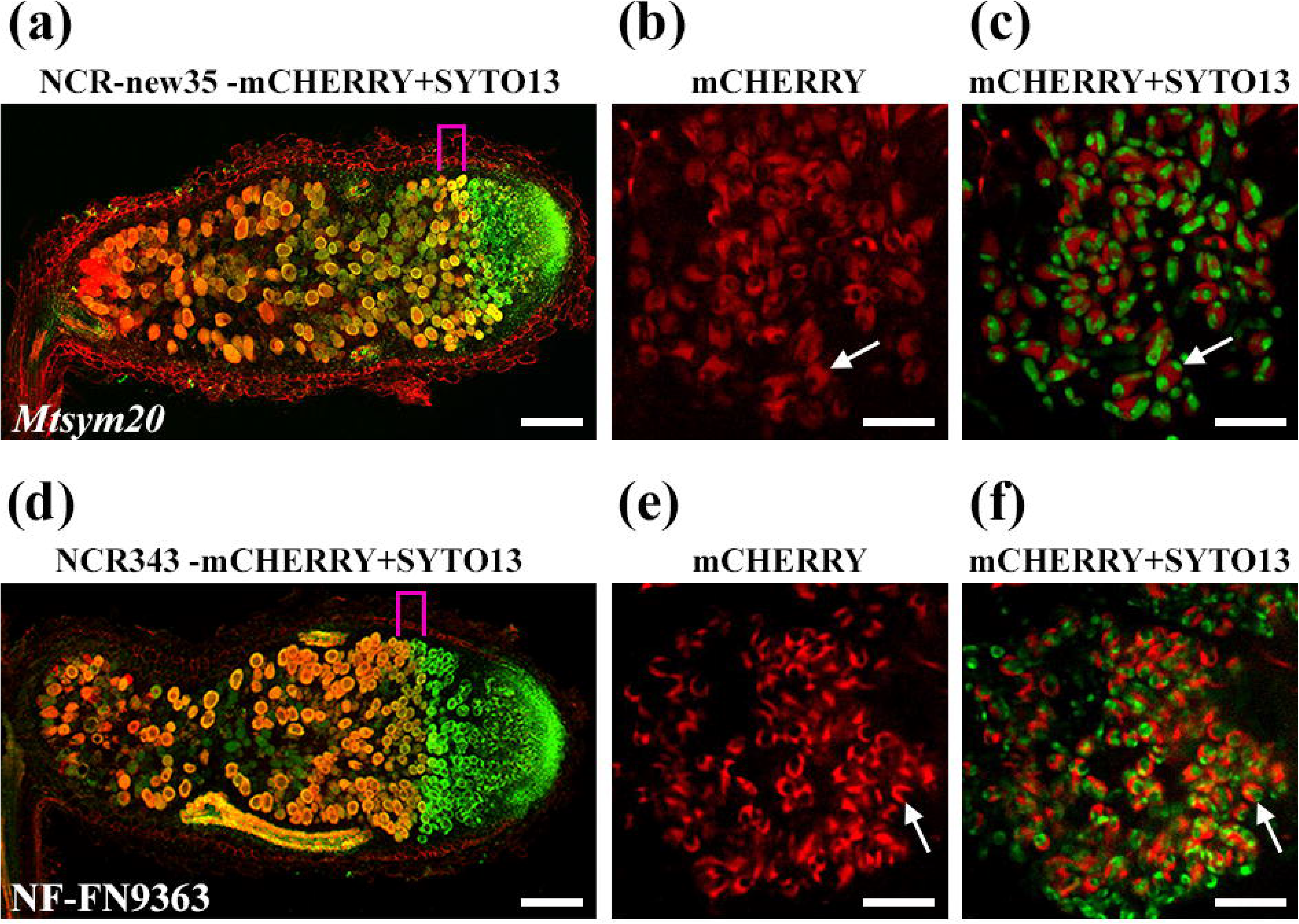
NCR-new35-mCherry and NCR343-mCherry fusion proteins restored the symbiotic defects of *Mtsym20* (a) and NF-FN9363 (d) mutant plants, respectively at 4 wpi with *S. medicae* WSM419. (a and d) The red NCR-new35-mCherry and NCR343-mCherry signals overlap with the green fluorescence of SYTO13-stained bacteroids in the infected cells of interzones (magenta brackets) and nitrogen fixation zones indicating the co-localization of bacteroids and peptides NCR-new35 and NCR343. (b, c, and e, f) Higher magnification of cells from the IZ revealed the red mCherry signal of NCR-new35 and NCR343 fusion proteins surrounding the bacteroids in the peribacteroid space. Bars, (a, d) 200 μm; (b, c, e and f) 20 μm.

## Discussion

The *M. truncatula* genome contains around 700 *NCR* genes which are almost exclusively expressed in symbiotic nodule cells (Roux et al. 2014). The encoded NCR peptides mediate terminal differentiation of the symbiotic partner (Van de Velde *et al*., 2010) but the high number of *NCR* genes raises the question of why *M. truncatula* evolved so many genes and how many of them are required for the establishment of symbiotic nitrogen fixation. Forward and reverse genetic analysis identified that peptides NCR169, NCR211 and NCR247 are crucial for effective nitrogen-fixing symbiosis (Horvath *et al*., 2015; Kim *et al*., 2015; Sankari *et al*., 2022) which prompted us to continue the analysis of ineffective *M. truncatula* symbiotic mutants to identify further essential *NCR* genes. Based on the nodulation phenotype of mutants defective in genes *NCR169*, *NCR211* and *NCR247*, we focused on symbiotic mutants *Mtsym19*, *Mtsym20* and NF-FN9363 that developed slightly elongated white nodules with zonation and invaded nodule cells in the distal part of the nodules. The histological analysis revealed that nodule zonation and the morphology of cells between the meristem and the transition zone in mutant nodules were very similar to wild-type nodules indicating the formation of differentiated symbiotic cell. However, the extended transition zone in mutant nodules compared with wt nodules indicated a defect in the differentiation process. This was in agreement with a former study detecting a higher ploidy level of nodule cells in *Mtsym19* and *Mtsym20* mutants compared with TE7 mutant, which is defective in the *IPD3* gene (Ovchinnikova *et al*., 2011; Horvath *et al*., 2011) with non-differentiated symbiotic nodule cells, but *Mtsym19* and *Mtsym20* nodule cells had lower endoreduplication index than wild type ones (Maunoury *et al*., 2010). The analysis of the length and DNA content of bacteroids isolated from *Mtsym19*, *Mtsym20* and NF-FN9363 nodules showed that mutant nodules contained lower proportion of longer and endoreduplicated bacteroids compared with wt nodules. These results indicated that the differentiation of nodule cells and hosted bacteroids was advanced but not complete in *Mtsym18*, *Mtsym19* and NF-FN9363 mutants. This finding was in accordance with the histological and transciptome analysis of *Mtsym19* and *Mtsym20* nodules which detected the activation of a second transcriptome-switch characteristic of late Fix mutant plants (Maunoury *et al*., 2010). The expression analysis of senescence and defence-related marker genes suggests that the lack of colonization of the cells in the mature part of nitrogen fixation zone and thus the absence of effective nitrogen fixation in mutant nodules induced premature senescence of symbiotic cells. The degradation of nodule cells and rhizobia was demonstrated in FN2106, another mutant deficient in *NCR343*, using a live/dead staining assay (Zhang et al. submitted). The transcript profiling analysis of FN2106 mutant also found the upregulation of senescence-related genes and no change or downregulation of genes associated with nodule defence response (Zhang et al. submitted).

The deletion in the NF-FN9363 mutant includes four *NCR* genes in close proximity. *NCR341*, *NCR343*, *NCR344* and *NCR345* showed high similarity at amino acid level which implies that these homologues have recently evolved at the scale of evolution by tandem duplication. However, only NCR343 is able to restore the symbiotic phenotype of NF-FN9363 indicating a specialization of this peptide for the symbiotic interaction between *M. truncatula* Jemalong A17 and the tested rhizobia. Further analysis by exchanging residues or fragments between NCR341, NCR343, NCR344 and NCR345 peptides could reveal the unique region(s) of NCR343 responsible for its distinct biological function. The identified deletions in *Mtsym19* and *Mtsym20* overlapped indicating that they are defective in the same gene which result is in contradiction to the previous allelism test that defined them as being in distinct complementation groups (Morandi *et al*., 2005). Our allelism test and complementation experiments with gene *NCR-new35* clearly confirmed that *Mtsym19* and *Mtsym20* are allelic. The spatial expression analysis of *NCR* genes identified by forward genetic approach and crucial for symbiosis (Horvath *et al*., 2015; Kim *et al*., 2015 and this study) was analysed by a GUS-reporter assay which showed that *NCR169*, *NCR211* and *NCR343* are active in the infected cells of IZ and ZIII but revealed that *NCR-new35* can be detected only in IZ cells. This observation is in correlation with RNA sequencing data obtained from laser-capture microdissected nodule zones (Roux *et al*., 2014). In addition to the expression of *NCR-new35* being detected predominantly only in IZ, its activity was much weaker compared with the other three *NCR* genes suggesting that the transcriptional intensity of an *NCR* gene does not correlate with the necessity of the *NCR* genes. Although the promoter activity of *NCR-new35* is barely detectable in nitrogen fixation zone, the encoded peptide is abundant in this zone suggesting a slow turnover or the enhanced stability of NCR-new35. Formerly, the identification of several NCR peptides, which are expressed during the early stages of nodule colonization, in mature bacteroids suggested the sustained stability of these NCR peptides (Durgo *et al*., 2015). Correspondingly to the previous finding on NCR169 peptide (Horvath *et al*., 2015), both NCR343 and NCR-new35 co-localise with bacteroids forming a ring-shaped fluorescent signal. Apparently, the fluorescent protein fusions of NCR peptides, which are functional and able to rescue the symbiotic phenotype of the corresponding mutants, are localised around the bacteroids. The co-localization of NCR343-YFP fusion protein with bacteroids has been also demonstrated in a related study (Zhang et al. submitted). However, a previous proteomic study identified NCR343 and NCR169, but not NCR211 and NCR-new35, in bacteroids (Durgo *et al*., 2015) indicating that at least a subset of the fluorescent-tagged NCR fusion molecules, conceivably following proteolytic cleavage, enter into the bacteroids.

The *NCR343* gene consists of one exon but *NCR-new35* has two exons, which is the most common gene structure of *M. truncatula NCR* genes. The encoded mature peptides are composed of 43 and 47 amino acids, respectively, including four cysteine residues at conserved positions, the characteristic feature of *M. truncatula* NCR peptides. The charge of the mature peptides is slightly and strongly anionic (isoeletric point (pI)=6.34 for NCR343 and 4.78 for NCR-new35). The charges of the five NCR peptides (NCR169 pI=8.45, NCR211 pI=5.38, NCR247 pI=10.15, NCR343 and NCR-new35), proved to be essential for symbiosis (Horvath *et al*., 2015; Kim *et al*., 2015; Sankari *et al*., 2022) vary between 4.78 and 10.15 indicating that anionic, neutral and cationic NCR peptides could be essential for nitrogen-fixing symbiosis in *M. truncatula*. The formation of intramolecular disulfide bonds between the conserved cysteine residues in NCR044 and NCR169 peptides was experimentally verified (Velivelli *et al*., 2020; Isozumi *et al*., 2021) and the *in planta* functional requirement of cysteine residues was demonstrated with substitution of any cysteine with serine which resulted in inactivation of NCR169 (Horvath *et al*., 2015). The replacement of the first cysteine residue with serine in NCR343 and NCR-new35 also abolished the symbiotic activity of the peptides implying that the requirement of cysteine residues for the *in planta* activity of NCRs is a common feature. Apart from the four cysteine residues in conserved position, the five crucial NCR peptides show high sequence variation but the presence of a valine, an isoleucine and one or two aspartic acid residues between the first two cysteines is dominant (Figure 4d). The relevance of these residues for the structure and symbiotic activity of NCR peptides requires further investigation. In brief, the only common feature among the five identified NCR peptides essential for symbiosis thus far (NCR169, NCR211, NCR247, NCR343 and NCR-new35) is that they belong to the subgroup of peptides with four conserved cysteine residues.

Based on the observation that terminally differentiated bacteroids are symbiotically more effective compared with reversibly differentiated ones (Oono *et al*., 2010), it is widely accepted that the controlled bacteroid growth in nodules provides increased fitness benefits for the host. In IRLC legumes and some Dalbergoid legume species, rhizobia undergo terminal differentiation provoked by NCR peptides (see reviews Pan & Wang, 2017; Downie & Kondorosi, 2021; Czernic *et al*., 2015). The large number of *NCR* genes of *M. truncatula* induced in consecutive steps and showing overlap expression which implies that these peptides might function together to optimise the interaction between rhizobia and the host, and potentially some of them compensate the negative effect of other NCR peptides (Mergaert, 2018). It has been presumed that there is a core set of *M. truncatula* NCRs with common functions, loss of which results in the termination of bacteroid differentiation and/or losing the viability of hosted rhizobia (Pan & Dong, 2017; Roy *et al*., 2020). Till now NCR169, NCR211 and NCR247 were identified as essential peptides for symbiosis but our work provides two additional NCR peptides, NCR343 and NCR-new35 required for effective nitrogen fixation between *M. truncatula* and *Sinorhizobium* species. The identification of novel crucial NCRs implies that further forward and reverse genetic studies might extend the cluster of universal and essential NCR peptides of *M. truncatula*.

## Acknowledgements

This work was supported by the Hungarian National Research Fund/National Research, Development and Innovation Office grants OTKA-67576, PD104334/108923, 106068, 119652 and 120122/120300, PD-121110 and PD-132495 as well as by the Collaborative Research Programme ICGEB Research Grant HUN17-03. The present work has benefited from Imagerie◻Gif core facility supported by l’Agence Nationale de la Recherche (ANR-11-EQPX-0029/Morphoscope, ANR-10-INBS-04/FranceBioImaging ; ANR◻11◻IDEX◻0003◻02/ Saclay Plant Sciences). We thank H. Cs. Tolnainé and K. Miró for their skillful technical assistance and A. Farkas for carrying out sample preparation and SEM imaging,

## Author contributions

B.H, Á.D., R.C. and P.K designed the project. B.H, B.G, M.T., Á.D, F.A., F.S., Y.C., M.B., J.B.B., and Z.T. performed the experiments, ÁD and Z.S. carried out data analysis. P.K wrote the manuscript.

## Data availability

All data supporting the findings of this study are available within the paper and within its supplementary materials published online.

---

**Fig. S1** Strain dependence of the nodulation and symbiotic phenotype of four ineffective symbiotic mutants NF-FN9363, *Mtsym20, Mtdnf4-1* and *Mtdnf7-2* compared with wt *M. truncatula* Jemalong nodules. (a) Plants were inoculated with *S. meliloti* strains 1021 and FSM-MA as well as with *S. medicae* strains ABS7 and WSM419 (*S. medicae* WSM419) and then nodule images were captured 3 wpi. Pink nodules were formed on wt roots with the four rhizobia strains. Roundish or slightly elongated white nodules developed on *Mtdnf7-2*, *Mtdnf4-1*, NF-FN9363 and *Mtsym20* roots, irrespective of the bacterial strains used for inoculation. Scale bars: 200μm. (b) Box-plot of nodule number and dry weight of wt and four ineffective symbiotic nitrogen-fixing mutants 3 weeks post inoculation with different rhizobia strains.

**Fig. S2** Transcriptional activity of senescence and defense-related marker genes in the nodules of four ineffective symbiotic mutants NF-FN9363, *Mtsym20*, *Mtdnf4-1, Mtdnf7-2* and *nad1-3* compared with wt nodules 3 wpi with *S. medicae* WSM419 using RT-qPCR. Senescence marker genes *MtCP2 and MtCP6* (a) were activated in all mutant nodules but pathogen-related genes *MtCHI* (b) and *MtPR10* (c) were highly activated only in *nad1-3* nodules.

**Fig. S3** Scanning electron microscopy analysis of symbiotic cells and bacteroid morphology in NF-FN9363, *Mtsym20*, *Mtdnf7-2*, *Mtdnf4-1* mutant and wt nodules 2 wpi with *S. medicae* WSM419. SEM images of whole nodule sections of wt plant (a1), mutant lines (b1-e1) free living *S. medicae* WSM419 bacteria (f1). Higher magnification of symbiotic cells characteristic for the distal part of infection zone (ZIId, a2-e2), the proximal part of infection zone (ZIIp, a3-e3), the first few cell layer of interzone (IZd, a4-e4), the proximal part of IZ of mutant nodules (IZp, b5-e5) and the mature nitrogen fixation zone (ZIII, a6-e6). An enlarged part of the symbiotic cells showing bacterial morphology are below each images. Scale bars: 100 μm in (a1-e1), 10 μm in (a2-e6). Scale bars in the images of plant cells and bacteroids at higher magnification (f1, a2-e6) 1 μm. it: infection thread b: bacteroid, db: decomposing bacteria, sb: saprophytic bacteria, v: vacuole, n: nucleus, s: starch granules *: discontinuity between rhizobia and plant cell walls.

**Fig. S4** Chromosomal position of the symbiotic loci of NF-FN9363, *Mtsym19* and *Mtsym20* and the allelism test between *Mtsym19* and *Mtsym20*. (a) The symbiotic locus of mutant NF-FN9363 was mapped in Linkage Group 6 (LG 6) of *M. truncatula* downwards the genetic marker Crs towards MtB178. Genetic mapping defined the chromosomal position of the symbiotic loci of *Mtsym20* and *Mtsym19* on chromosome 4 (LG 4). (b) The allelic relationship of *Mtsym20* and *Mtsym19* mutants was proved with allelism tests. F3 *Mtsym20* and *Mtsym19* symbiotic mutant plants, selected from the mapping populations showing opposite parental homozygous genotypes for the genetic marker MtB267, were crossed and the offsprings were scored for the symbiotic phenotype. The success of the crossing of the two mutant lines was detected with the heterozygous genotypes of C1, C3 and C4 hybrid plants. Plants of C2 and C5 were self-mated. The Fix- symbiotic phenotype of plants of C1, C3 and C4 crosses revealed the allelic relationship of mutants *Mtsym20* and *Mtsym19*; MM, Molecular weight Marker

**Fig. S5** The deleted genomic region in the symbiotic locus of NF-FN9363. (a) An approximately 500 kb genomic region deleted in the symbiotic locus in NF-FN9363 was defined with PCR-based markers designed for genes. The A17 genomic DNA was used as a wild type control in the PCR reactions. * primer dimers. M, mutant-NF-FN9363; MM, Molecular weight Marker (b) List of genes or gene models in the deleted genomic region detected by using microarray-based CGH. Dark grey shading shows genes corresponding to probesets with log2 CGH ratios lower than −2,5, pale grey shading is for log_2_ CGH ratios less than −1 and greater than −2,5. (+) and (−) indicate the presence or absence of PCR fragments identified with the genomic DNA of NF-FN9363. Deleted *NCR* genes in the symbiotic locus of NF-FN9363 are in bold.

**Fig. S6** The deleted genomic region of the symbiotic loci of *Mtsym19* and *Mtsym20*. (a) PCR-based markers identified overlapping deleted genomic regions in *Mtsym19* and *Mtsym20* on chromosome 4 (Chr 4). PCR-based markers were designed for genes marked with gene IDs and visualized on agarose gel. The *M. truncatula* Jemalong (J5) genomic DNA was used for a wild-type control. (b) List of genes and gene models in the deleted genomic region. (+) not deleted: (−) deleted genes in the symbiotic loci of *Mtsym19* and *Mtsym20* detected with PCR-based markers. MM, Molecular weight Marker

**Fig. S7** Complementation assays of NF-FN9363, *Mtsym20* and *Mtsym19* symbiotic ineffective mutants with *NCR* gene constructs and the study of the requirement of first cysteine residues in the function for the peptides NCR343, NCR-new35 and NCR211. Constructs of *NCR* genes controlled by their native promoter fragments were introduced into the roots of mutant plants using *A. rhizogenes*-mediated transformation. The transgenic roots were detected either by dsRed fluorescence, in the case of gene constructs created in single site gateway (GW) system (a1-i1), or with GFP fluorescence using the constructs generated by multi-site GW system (j1-t1). Complementation was assessed by the formation of cylindrical nodules with the characteristic zonation of indeterminate nodules. To compare zonation of nodules, nodules developed on the wild-type (a1-a3 and j1-j3) and symbiotic mutant roots (*Mtsym20* - c1-c2 and p1-p3, *Mtsym19* - e1-e2, NF-FN9363 - g1-g3 and l1-l3, and *Mtdnf4-1*-s1-s3) transformed with empty vector were used. The symbiotic defects of both mutants *Mtsym20* (b1-b3 and o1-o3) and *Mtsym19* (d1-d3) were rescued with the *NCR-new35* gene constructs indicating that they are deficient in the same genes. The *NCR343* gene constructs (f1-f3 and k1-k3) restored but the constructs of *NCR345* (h1-h3), *NCR341* (i1-i3) and *NCR344* (m1-m3) did not complement the nodulation phenotype of NF-FN-9363. The modification of the constructs substituting the first cysteines for serines equally abolished the ability of NCR343-C34S (n1-n3), NCR-new35-C41S (q1-q3), NCR211-C28S (t1-t3) peptides to restore the symbiotic nodulation phenotype of mutants NF-FN9363, *Mtsym20* and *Mtdnf4-1*, respectively. Construct containing the wild-type *NCR211* gene restored the symbiotic nodulation phenotype on *Mtdnf4-1* roots (r1-r3). Bars, (a1-t3) 200 μm.

