## Supplementary material for "The *Medicago truncatula* nodule-specific cysteine-rich peptides, NCR343 and NCR-new35 are required for the maintenance of rhizobia in nitrogen-fixing nodules"

**Figure S1**

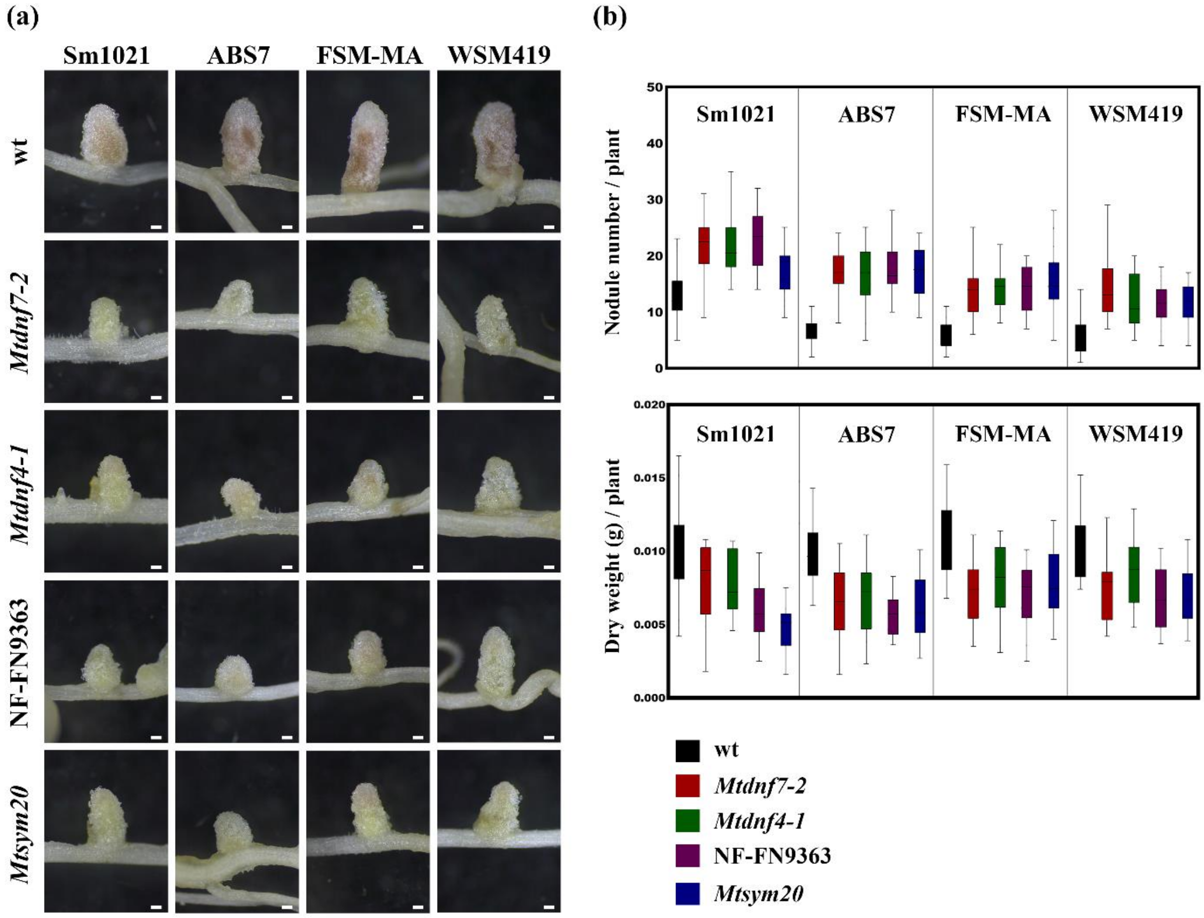
(a)
Sm1021
ABS7
FSM-MA
WSM419
wt
Mtdnf7-2
Mtdnf4-1
NF-FN9363
Mtsym20
(b)
Nodule number / plant
Dry weight (g) / plant
Sm1021
ABS7
FSM-MA
WSM419
wt
Mtdnf7-2
Mtdnf4-1
NF-FN9363
Mtsym20

#### Figure S2

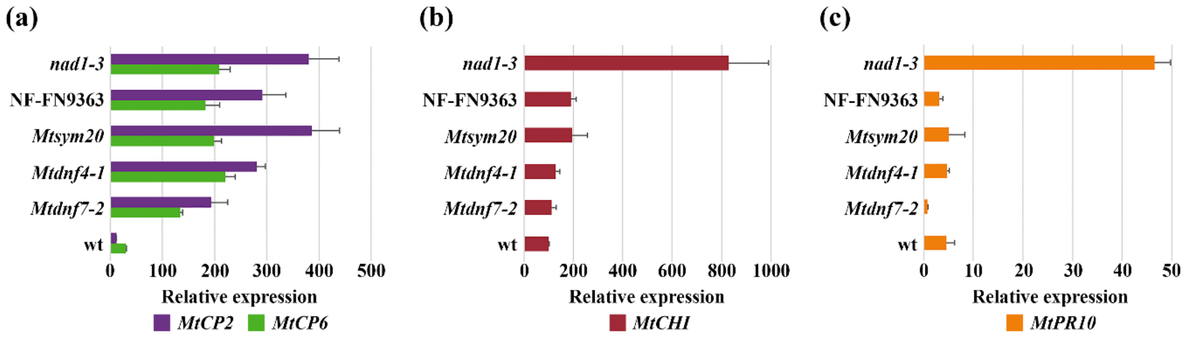

(a)
nad1-3
NF-FN9363
Mtsym20
Mtdnf4-1
Mtdnf7-2
wt
0 100 200 300 400 500
Relative expression
MtCP2 MtCP6
(b)
nad1-3
NF-FN9363
Mtsym20
Mtdnf4-1
Mtdnf7-2
wt
0 200 400 600 800 1000
Relative expression
MtCHI
(c)
nad1-3
NF-FN9363
Mtsym20
Mtdnf4-1
Mtdnf7-2
wt
0 10 20 30 40 50
Relative expression
MtPR10

**Figure S3**

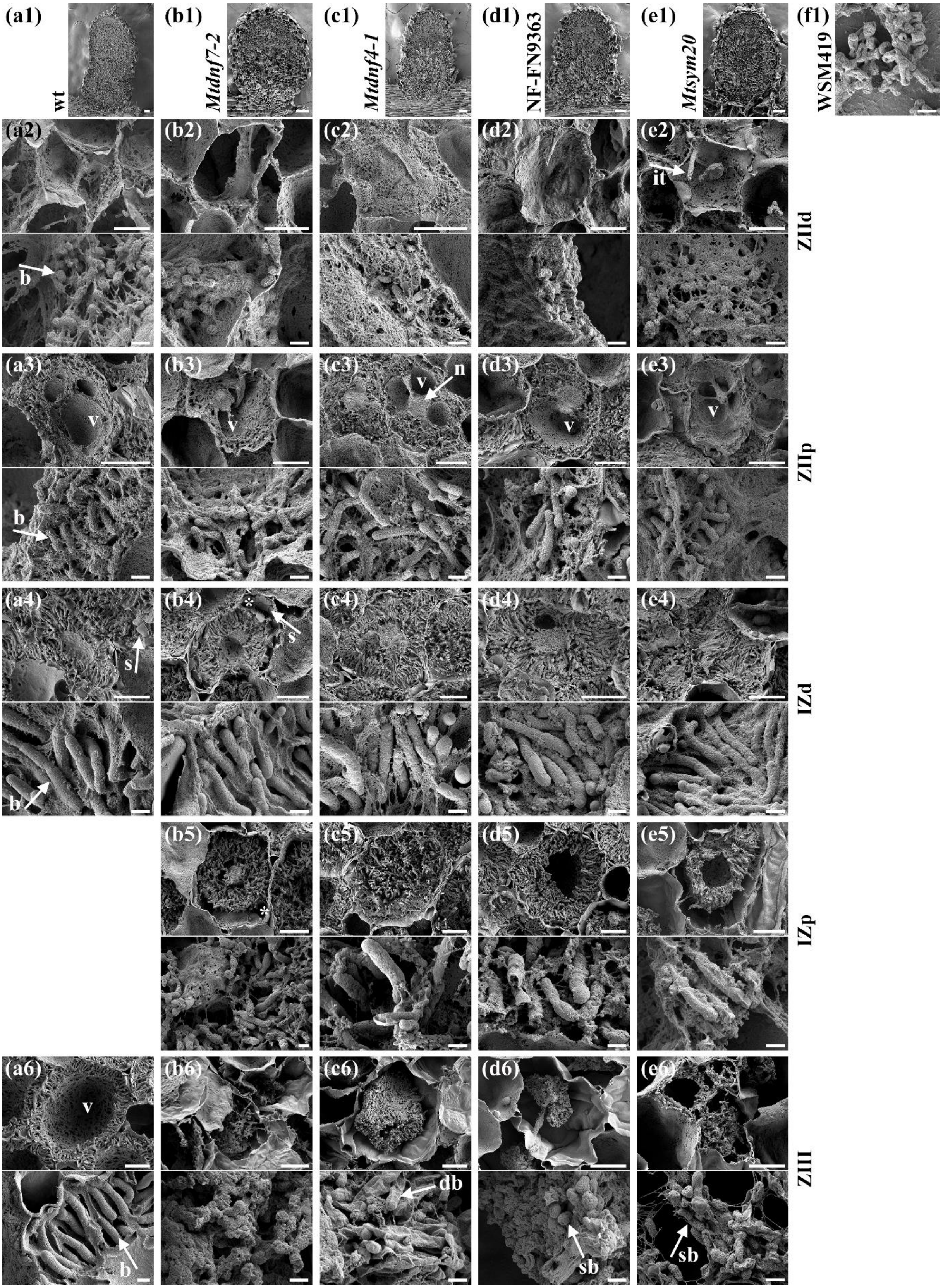

(a1)
wt
(b1)
Mtdnf7-2
(c1)
Mtdnf4-1
(d1)
NF-FN9363
(e1)
Mtsym20
(f1)
WSM419
(a2)
(b2)
(c2)
(d2)
(e2)
it
b
ZIId
(a3)
(b3)
(c3)
(d3)
(e3)
v
n
b
ZIIp
(a4)
(b4)
(c4)
(d4)
(e4)
s
*
b
IZd
(b5)
(c5)
(d5)
(e5)
IZp
(a6)
(b6)
(c6)
(d6)
(e6)
db
sb
ZIII

#### Figure S4

(a)

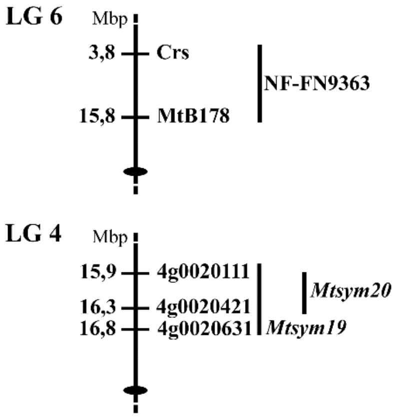


(b)

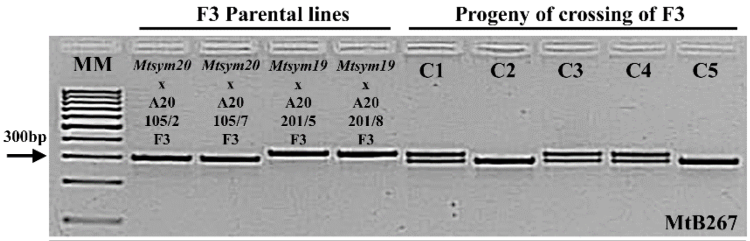


| Progeny of crossing of F3 plants | C1 | C2 | C3 | C4 | C5 |
|---|---|---|---|---|---|
| Crosses | *Mtsym20* xA20 105/2_F3 X *Mtsym19* xA20 201/8_F3 | *Mtsym20* xA20 105/2 _F3 X *Mtsym19* xA20 201/5_F3 | *Mtsym20* xA20 105/2_F3 X *Mtsym19* xA20 201/8_F3 | *Mtsym20* xA20 105/7_F3 X *Mtsym19* xA20 201/8_F3 | *Mtsym20* xA20 105/7_F3 X *Mtsym19* xA20 201/5_F3 |
| Number of progeny | 4 | 4 | 1 | 3 | 4 |
| Phenotype | Fix- | Fix- | Fix- | Fix- | Fix- |

### Figure S5

**(a)**

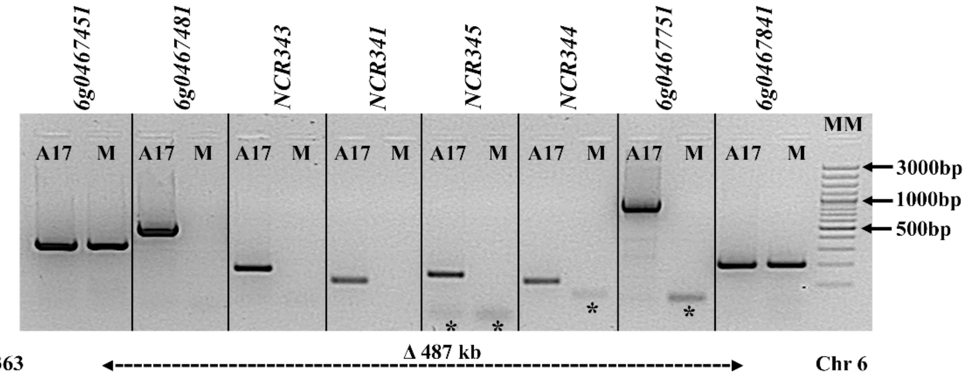


**(b)**

| Gene ID IMGA-Mt3.5.1 | Gene ID JCVI-Mt4.0-mRNA | Gene ID MtrunA17r 5.0-ANR | Location | Description | PCR test NF-FN9363 | CGH Probasets | Log2 Ratios NF-FN9363/A17 |
|---|---|---|---|---|---|---|---|
| Medtr6g044320 6;8584340;8584714 | Medtr6g044320.1 | MtrunA17Chr6g0467451 | 16368812..16369398 | putative protein | + | CHR06FS008584352 | (-0,658277) |
| Medtr6g044390; 6:8608190:8610027 | Medtr6g044390.1-2 | MtrunA17Chr6g0467461 | 16392692..16394407 | Putative major intrinsic protein | | CHR06FS008608662, CHR06FS008608762, CHR06FS008608862, CHR06FS008609062, CHR06FS008609162, CHR06FS008609262, CHR06FS008609462, CHR06FS008609662, CHR06FS008609762, CHR06FS008611262, CHR06FS008611462 | (-0,0555089)-(-0,80517) |
| Medtr6g044400 6;8611053;8612217 | Medtr6g044400.1 | MtrunA17Chr6g0467471 | 16395439..16398804 | hypothetical protein | - | CHR06FS008612462, CHR06FS008612862 | (-4,55433)-(-4,88239) |
| Medtr6g044410; 6;8612281;8615924 | Medtr6g044410.1 | MtrunA17Chr6g0467481 | 16396749..16397894 | putative protein | | | |
| Medtr6g044470 6;8630183;8632239 | Medtr3g024060.1 | MtrunA17Chr6g0467491 | 16415208..16416884 | putative protein | | CHR06FS008630188, CHR06FS008630388, CHR06FS008631988, CHR06FS008632088 | (-0,170679)-(-0,955089) |
| Medtr6g044560 6;8665344;8668128 | Medtr6g044560.1<br>Medtr6g044572.1 | MtrunA17Chr6g0467501 | 16449684..16452467 | Putative F-box domain, FBD domain, plant specific mitochondrial import receptor subunit TOM20 | | CHR06FS008665365, CHR06FS008667165, CHR06FS008667365 | (-2,69474)-(-3,95663) |
| Medtr6g044570 6:8668454:8673396 | Medtr6g044562.1<br>Medtr6g044575.1 | MtrunA17Chr6g0467511 | 16452075..16453323 | **NCR343 Nodule-specific Cysteine-Rich peptide** | - | CHR06FS008668574, CHR06FS008668774, CHR06FS008668874, CHR06FS008668974, CHR06FS008669174 | (-0,855741)-(-1,52003) |
| Medtr6g044600 6:8693298:8699693 | Medtr6g044600.1 | MtrunA17Chr6g0467521 | 16477585..16484099 | Putative chromatin regulator PHD family | | CHR06FS008693381, CHR06FS008694539, CHR06FS008695439, CHR06FS008695539, CHR06FS008696039, CHR06FS008697939 | (-2,52586)-(-2,7707) |
| Medtr6g044690 6:8731563;8735699 | Medtr6g044690.1 | MtrunA17Chr6g0467531 | 16515897..16517600 | Putative [Histone H3]-lysine(4) N-trimethyltransferase chromatin remodeling SET family | | CHR06FS008731665, CHR06FS008732065, CHR06FS008732165, CHR06FS008732465, CHR06FS008732565, CHR06FS008732665, CHR06FS008732865, CHR06FS008732965, CHR06FS008733165, CHR06FS008733365, CHR06FS008733465, CHR06FS008733565, CHR06FS008733765, CHR06FS008733865, CHR06FS008733965, CHR06FS008734165, CHR06FS008734265, CHR06FS008734465, CHR06FS008734665, CHR06FS008734965, CHR06FS008735565 | 0,675527-(-1,24204) |
| | no annotated gene | MtrunA17Chr6g0467541 | 16518840..16519902 | hypothetical protein | | no probes in this region | |
| Medtr6g044700 6:8736003:8736554 | Medtr6g044700.1 | MtrunA17Chr6g0467551 | 16520361..16521072 | **NCR341 Nodule-specific Cysteine-Rich peptide** | - | CHR06FS008736066, CHR06FS008736166, CHR06FS008736366, CHR06FS008736466 | (-0,0798214)-(-1,54029) |
| Medtr6tc044720 6:8740565:8741683 | Medtr6tc044720.1 | MtrunA17Chr6g0467561 | 16524899..16525180 | Putative RNA-directed DNA polymerase | | no probes in this region | |
| Medtr6g044730 6;8742038;8747814 | Medtr6g044735.1 | MtrunA17Chr6g0467571 | 16526347..16527313 | **NCR345 Nodule-specific Cysteine-Rich peptide** | - | CHR06FS008742867 | (-3,29496) |
| | Medtr6g044730.1 | MtrunA17Chr6g0467581 | 16531872..16532960 | **NCR344 Nodule-specific Cysteine-Rich peptide** | - | CHR06FS008747671 | (-4,28487) |
| Medtr6g044740 6:8749719:8753098 | Medtr6g044740.1 | MtrunA17Chr6g0467591 | 16533627..16534375 | putative protein | | CHR06FS008749771, CHR06FS008749871, CHR06FS008750271, CHR06FS008750371, CHR06FS008750471, CHR06FS008750671, CHR06FS008750771, CHR06FS008750871, CHR06FS008751071, CHR06FS008751171, CHR06FS008751271, CHR06FS008751571, CHR06FS008751871, CHR06FS008752071, CHR06FS008752371, CHR06FS008752871, CHR06FS008753071 | (-0,716866)-(-2,48381) |
| | | MtrunA17Chr6g0467601 | 16534377..16537693 | Putative protein kinase RLK-Pelle-WAK-LRK10L-1 family | | CHR06FS008751471, CHR06FS008751671, CHR06FS008751971, CHR06FS008752271 | (-2,53658)-(-3,75131) |

| IMGA-Mt3.5.1 | Gene ID JCVI-Mt4.0-mRNA | MtrunA17r 5.0-ANR | Location | Description | PCR test NF-FN9363 | CGH Probasets | Log2 Ratios NF-FN9363/A17 |
|---|---|---|---|---|---|---|---|
| Medtr6g044780 6:8761825:8764598 | Medtr6g044780.1 | MtrunA17Chr6g0467611 | 16546202..16547182 | Kunitz-P Trypsin Inhibitor | | CHR06FS008762304, CHR06FS008762604, CHR06FS008762704, CHR06FS008762904, CHR06FS008763004, CHR06FS008763104, CHR06FS008763304, CHR06FS008763404, CHR06FS008763504, CHR06FS008763704, CHR06FS008763804, CHR06FS008763904, CHR06FS008764104, CHR06FS008764204, CHR06FS008764304, CHR06FS008764504 | (-0,943527)-(-2,45445) |
| Medtr6g044810 6:8769009:8769958 | Medtr6g044810.1 | MtrunA17Chr6g0467621 | 16553380..16554337 | Kunitz-P Trypsin Inhibitor | | CHR06FS008769304, CHR06FS008769904 | (-1,19029)-(-1,58464) |
| Medtr6g044830: 6:8776617:8780252 | Medtr6g044830.1 | MtrunA17Chr6g0467631 | 16560875..16564797 | Putative protein kinase RLK-Pelle-WAK-LRK10L-1 family | | CHR06FS008776704, CHR06FS008776804, CHR06FS008777804, CHR06FS008778104, CHR06FS008779604, CHR06FS008779804 | (-2,68171)-(-4,19058) |
| Medtr6g044960 6:8829979:8832047 | Medtr6g445600.1<br>Medtr0690s0010.1 | MtrunA17Chr6g0467641 | 16608614..16610573 | Putative glycerophosphodiester phosphodiesterase, protein kinase RLK-Pelle-LRK10L-2 family | | no probes in this region | |
| Medtr6g045080 6:8873343:8873737 | Medtr6g445600.1<br>Medtr0690s0010.1 | MtrunA17Chr6g0467651 | 16611148..16611498 | Putative glycerophosphodiester phosphodiesterase, protein kinase RLK-Pelle-LRK10L-2 family | | CHR06FS008873513 | (-1,67102) |
| Medtr6g044960: 6:8829979:8832047<br>Medtr6g045090: 6:8874098:8881712 | Medtr6g445600.1<br>Medtr0690s0010.1 | MtrunA17Chr6g0467661 | 16614494..16616544 | Putative glycerophosphodiester phosphodiesterase, protein kinase RLK-Pelle-LRK10L-2 family | | CHR06FS008876313, CHR06FS008878303, CHR06FS008878403, CHR06FS008878503, CHR06FS008878703, CHR06FS008878803, CHR06FS008878903, CHR06FS008879103, CHR06FS008879203, CHR06FS008879503, CHR06FS008879603, CHR06FS008879703, CHR06FS008879903, CHR06FS008880003, CHR06FS008880103, CHR06FS008880303, CHR06FS008880403, CHR06FS008880503, CHR06FS008880703, CHR06FS008880803, CHR06FS008880903, CHR06FS008881103, CHR06FS008881203, CHR06FS008881303, CHR06FS008881503, CHR06FS008881703 | 0,661857-(-1,36335) |
| Medtr6g044960: 6:8829979:8832047 | Medtr6g445600.1<br>Medtr0690s0010.1 | MtrunA17Chr6g0467671 | 16617103..16617453 | Putative glycerophosphodiester phosphodiesterase, protein kinase RLK-Pelle-LRK10L-2 family | | no probes in this region | |
| Medtr6g044960: 6:8829979:8832047 | Medtr6g445600.1<br>Medtr0690s0010.1 | MtrunA17Chr6g0467681 | 16620561..16622518 | Putative glycerophosphodiester phosphodiesterase, protein kinase RLK-Pelle-LRK10L-2 family | | no probes in this region | |
| Medtr6g044960: 6:8829979:8832047 | Medtr0690s0010.1 | MtrunA17Chr6g0467691 | 16623060..16623410 | Putative glycerophosphodiester phosphodiesterase, protein kinase RLK-Pelle-LRK10L-2 family | | no probes in this region | |
| Medtr6g044960: 6:8829979:8832047 | Medtr0690s0010.1 | MtrunA17Chr6g0467701 | 16626577..16627853 | Putative glycerophosphodiester phosphodiesterase, protein kinase RLK-Pelle-LRK10L-2 family | | no probes in this region | |
| Medtr6g044970 6:8832719:8833673<br>Medtr6g044980 6:8836146:8836572<br>Medtr6g044990 6:8838885:8839677<br>Medtr6g045000: 6:8840297:8840751<br>Medtr6g045010 6:8840936:8842077<br>Medtr6te045020 6:8843697:8845661<br>Medtr6g045030 6:8846480:8847885 | Medtr6g044970.1<br>no annotated gene<br>no annotated gene<br>no annotated gene<br>Medtr6g045010.1<br>Medtr6te045025.1<br>Medtr6g045030.1 | MtrunA17Chr6g0467711 | 16629255..16644713 | Putative glycerophosphodiester phosphodiesterase, protein kinase RLK-Pelle-LRK10L-2 family | | CHR06FS008833533, CHR06FS008839062, CHR06FS008841562, CHR06FS008841762, CHR06FS008841962, CHR06FS008842062 | (-2,58058)-(-4,1295) |
| no annotated gene | Medtr6g045057.1 | MtrunA17Chr6g0467721 | 16668911..16669662 | Putative potassium transporter | | no probes in this region | |
| no annotated gene | no annotated gene | MtrunA17Chr6g0467731 | 16704349..16704591 | hypothetical protein | | no probes in this region | |
| no annotated gene | Medtr6g045077.1 | MtrunA17Chr6g0467741 | 16708723..16709781 | Putative Hemopexin-like domain-containing protein | | no probes in this region | |
| no annotated gene | Medtr6g045087.1 | MtrunA17Chr6g0467751 | 16715695..16719024 | Putative solute-binding protein family 3/ domain of MltF | - | no probes in this region | |
| no annotated gene | Medtr6g045097.1 | MtrunA17Chr6g0467761 | 16728979..16729889 | Kunitz-P Trypsin Inhibitor | | no probes in this region | |
| no annotated gene | Medtr6g045403.1 | MtrunA17Chr6g0467771 | 16770047..16770821 | Kunitz-P Trypsin Inhibitor | | no probes in this region | |
| no annotated gene | Medtr6g045433.1 | MtrunA17Chr6g0467781 | 16791349..16791917 | Kunitz-P Trypsin Inhibitor | | no probes in this region | |
| no annotated gene | no annotated gene | MtrunA17Chr6g0467801 | 16806306..16806986 | hypothetical protein | | no probes in this region | |
| no annotated gene | Medtr6g045467.1-6 | MtrunA17Chr6g0467811 | 16828529..16833926 | Putative gliding motility-associated protein GldF | | no probes in this region | |
| no annotated gene | no annotated gene | MtrunA17Chr6g0467821 | 16836080..16836189 | MIR2587 | | no probes in this region | |
| no annotated gene | Medtr6g045483.1-7 | MtrunA17Chr6g0467831 | 16840379..16843986 | Putative citrate transporter-like domain-containing protein | | no probes in this region | |
| no annotated gene | Medtr6g045493.1 | MtrunA17Chr6g0467841<br>MtrunA17Chr6g0467851 | 16851963..16856135<br>16852310..16856126 | Putative uncharacterized protein<br>Putative protein | + | no probes in this region | |

**Figure S6**

**(a)**

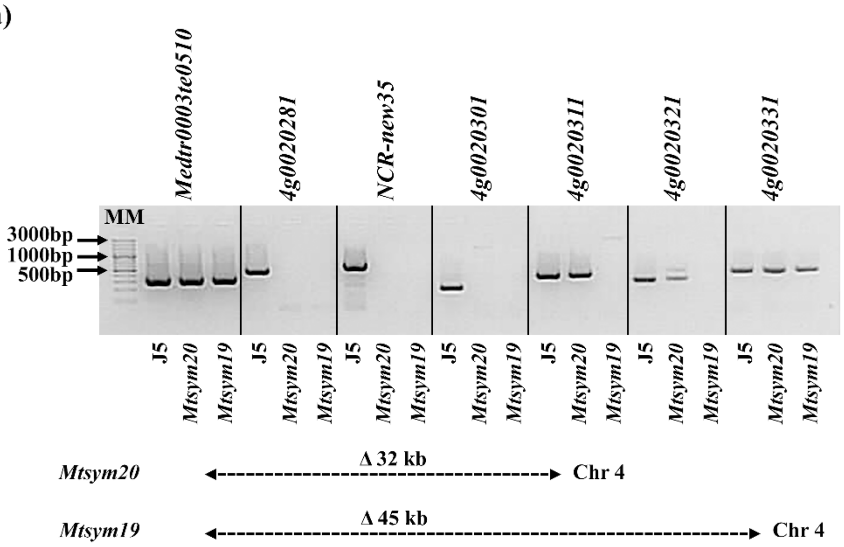


**(b)**

| Gene ID MtrunA17r5.0-ANR | Location | Description | PCR test *Mtsym20* | *Mtsym19* |
|---|---|---|---|---|
| Medtr0003te0510.1 | 16245316..16247149 | - | + | + |
| MtrunA17Chr4g0020281 | 16256649..16262089 | Putative protein | - | - |
| MtrunA17Chr4g0020291 | 16267079..16268349 | **Nodule-specific Cysteine-Rich peptide NCR-new35** | - | - |
| MtrunA17Chr4g0020301 | 16268538..16269391 | Putative protein | - | - |
| MtrunA17Chr4g0020311 | 16277110..16278540 | Putative long-chain-alcohol O-fatty-acyltransferase | + | - |
| MtrunA17Chr4g0020321 | 16279358..16287505 | Putative GIY-YIG endonuclease | + | - |
| MtrunA17Chr4g0020331 | 16290767..16294177 | Putative equilibrative nucleoside transporter | + | + |

**Figure S7**

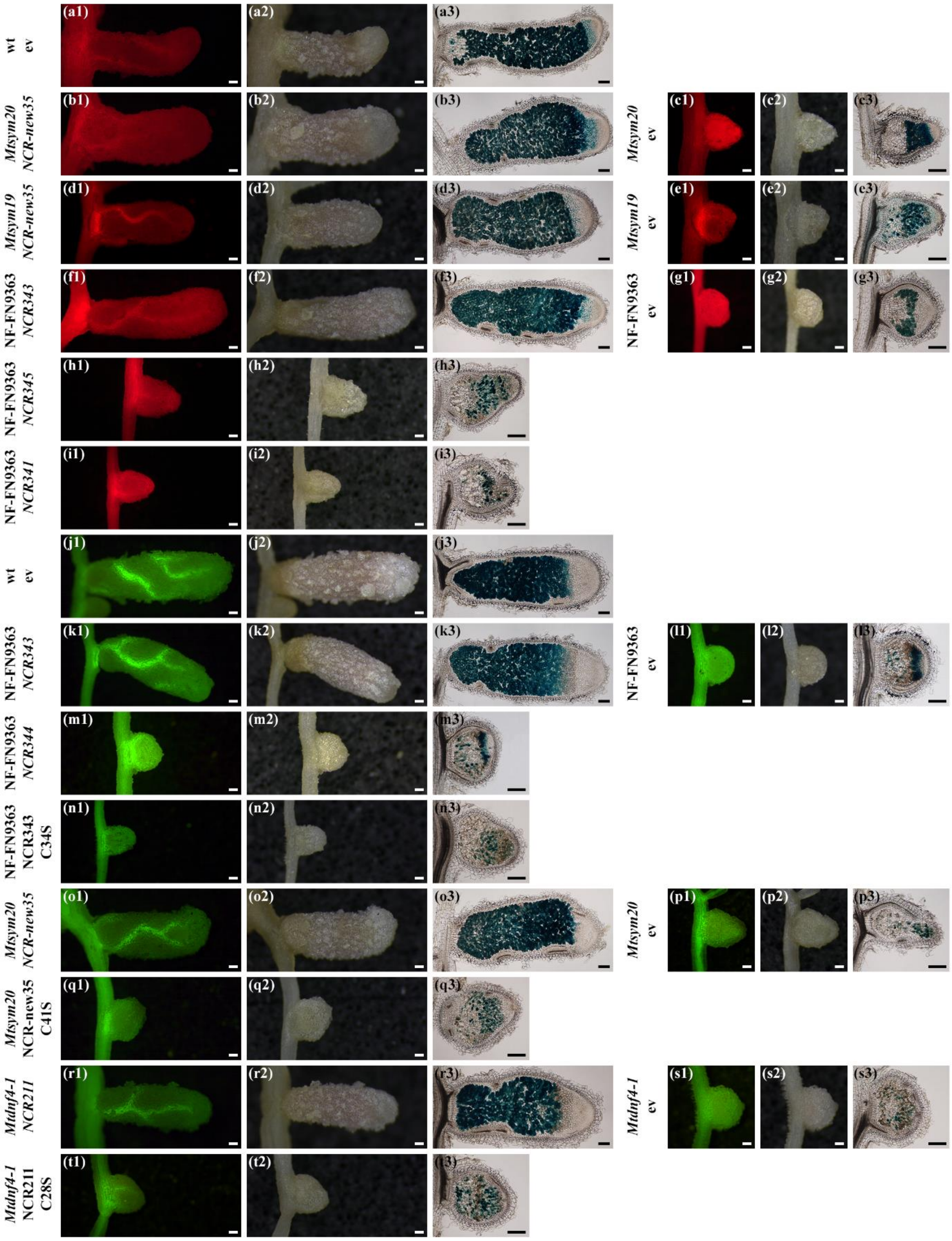

wt ev
Mtsym20 NCR-new35
Mtsym20 ev
Mtsym19 NCR-new35
Mtsym19 ev
NF-FN9363 NCR343
NF-FN9363 ev
NF-FN9363 NCR345
NF-FN9363 NCR341
wt ev
NF-FN9363 NCR343
NF-FN9363 ev
NF-FN9363 NCR344
NF-FN9363 NCR343 C34S
Mtsym20 NCR-new35
Mtsym20 ev
Mtsym20 NCR-new35 C41S
Mtdnf4-1 NCR211
Mtdnf4-1 ev
Mtdnf4-1 NCR211 C28S

**Table S1.** List of primers used to define the deletion borders in symbiotic mutant lines, generate gene constructs and analyse gene expression using RT-qPCR

| **Primers to define the deletion borders in symbiotic mutant lines:** | **Forward (5'→3')** | **Reverse (5'→3')** |
|---|---|---|
| **NF-FN9363:** | | |
| 6g0467451 | ACCGTAAACATATGTAGCTGTCA | CGTTTGCCATGAGTTTTGCC |
| 6g0467481 | AGAAACATGAGTCTGAAGGAGC | ATTGTTTAAGGGCCTGGTTTCC |
| 6g0467501 | AGATGGGGTGTTGCTTTGTTA | CGTCGTGCGAGTCTGGTGTTA |
| NCR343 | TCATTTCTTTCCATGTTTCTTGT | TGGCTTTTTGGGTATCTCTCTTA |
| NCR341 | TAAGTTTGTTTATGGTTTAGTTT | AGCATCAGTATCACATTTAATAT |
| NCR345 | TTCAAGTTTATTTATGGTTTAGTT | ATTGTCTACATAGTTTGTCAATGC |
| NCR344 | TTTTTATGCGGTTTAGTTTTATTT | CATTTTAGGACAACCAGCATCAGT |
| 6g0467751 | GTTTGGCTTCTATTGGCGTTTG | GTTTCTGTAGATGGTGTGGAGC |
| 6g0467841 | AAACCACACCACATGAGTCA | GTTGTTGTTGAGGCTACCCA |
| ***Mtsym19, Mtsym20:*** | | |
| Medtr0003te0510 | GACAGGTGGCTTTGGAGATT | CATTAGGGATTACCGAGCAGAC |
| 4g0020281 | TCCAAACGGAGCCTTAATCT | GCAAAATAGTGAAAGCCCGA |
| NCR-new35 | CAAATTTACAAGGACGTTTTGCA | AGGATACACGCATAAGAGACATTTA |
| 4g0020301 | AGGTATCTTTCGCCCAGA | CGGGGTGTTACATAATACATATAC |
| 4g0020311 | TGGTGACAAGCATCCTAAGAC | AAGCCCTAACATCAAGCCTAC |
| 4g0020321 | GGCGAGGGAAATTGGTAGTG | TCAAGAGCTGGTCGGAGAA |
| 4g0020331 | CAACAATGGTATTCACCGATTCC | CGACCCGACCCTTAACATAAA |
| **Primers used for RT-qPCR:** | **Forward (5'→3')** | **Reverse (5'→3')** |
| **Reference genes:** | | |
| 3g0126781(UBI)-q | GGCCCTAGAACATTTCCTGTGG | CAGTCTTCAAAACTCTTGGGCAG |
| 3g0126461(PTB)-q | CGCCTTGTCAGCATTGATGTC | TGAACCAGTGCCTGGAATCCT |
| **Detection of gDNA contamination:** | | |
| 3g0126781(UBI)-intron-q | GTCCTCTAAGGTTTAATGAACCGG | GAAAGACACAGCCAAGTTGCAC |
| **Expression analysis of *NCR* genes:** | | |
| NCR343-q | AGAGGTTGATGGTGCCTTTG | AAATTTCACATAGCTCGTTAACACAC |
| NCR014-q | TGATTGGAAGGAAAGGAGGACC | TGTGCAAAAGTTATTAACGCATAGC |
| NCR169-q | GGAAATGCGTTGAAAATGTTTGTG | AACATTTCTCCACTTTATTCTCGGG |
| NCR211-q | GCGTTAATGCACATCATTTGTGG | TTATTCTCGGACACAAACACCTTG |
| **Realitve expression of senescence and defence response marker genes :** | | |
| 4g0041081(CP6)-q | GCAATGATGGCATCTTATCCC | TACACAAGCATAATCACAATTTAC |
| 5g0405751(CP2)-q | CATCTTACCCTACTGCTTAAATGC | AACTAGAAACCATGATGAATGTAGC |
| 3g0145981(CHI)-q | GGGCTTGAATGCGGAAGAGG | CAAGATTGTCTCCATATCCAACTCC |
| 2g0295064(PR10)-q | TGTTGAAGATGGTGAGACCAAGC | GTCTGGAAGGCCAACACCTCC |
| **Primers used to generate gene constructs:** | **Forward (5'→3')** | **Reverse (5'→3')** |
| **Single site gateway (GW) clones in pKGW-prUBQ10::dsRed vector for complementation tests:** | | |
| NCR343-GW | GGGGACAAGTTTGTACAAAAAAGCAGGCTGGCAACATTTTTCTCTCTTATGCCTTT | GGGGACCACTTTGTACAAGAAAGCTGGGTGCGGTAGCGTTTCTCATTAGTTTCTA |
| NCR341-GW | GGGGACAAGTTTGTACAAAAAAGCAGGCTGGTATTTGTTCAAGTGTGCAGGGTC | GGGGACCACTTTGTACAAGAAAGCTGGGTGATTTTACGGCATGTGATTAGGGA |
| NCR345-GW | GGGGACAAGTTTGTACAAAAAAGCAGGCTGGTAGGTTGTTTCGTTGTTCTTC | GGGGACCACTTTGTACAAGAAAGCTGGGTGCAGTTATTGGCTACATTCGTT |
| NCR-new35-GW | GGGGACAAGTTTGTACAAAAAAGCAGGCTTGTTGTAGAAGGAAGAGGAGGAG | GGGGACCACTTTGTACAAGAAAGCTGGGTCCGTTTTGACAGCTCTAACAGTA |
| **Multisite gateway (MsGW) clones in pKm43GW-rolD::EGFP vector for complementation tests:** | | |
| pNCR344-MsGW | GGGGACAACTTTGTATAGAAAAGTTGGGCTTTAATATGTTGGATCCTATTCA | GGGGACTGCTTTTTTGTACAAACTTGGATTATTTTTTCTCGTTTCTTGTAA |
| NCR344-coding-MsGW | GGGGACAAGTTTGTACAAAAAAGCAGGCTGGATGGCTAAAACTCTCAAATTTTTA | GGGGACCACTTTGTACAAGAAAGCTGGGTGTTATGACCTTCTAAATTGTTTACA |
| NCR344-3UTR-MsGW | GGGGACAGCTTTCTTGTACAAAGTGGGGCATAAATCTCATAATATGTAGTTC | GGGGACAACTTTGTATAATAAAGTTGGTGGTGAACATAAGCTGCTGCTAAT |
| pNCR343-MsGW | GGGGACAACTTTGTATAGAAAAGTTGGGCAACATTTTTCTCTCTTATGCCTTT | GGGGACTGCTTTTTTGTACAAACTTGGATTTGTTTTCTTTGTTTTTGTAACA |
| NCR343-coding-MsGW | GGGGACAAGTTTGTACAAAAAAGCAGGCTGGATGGCTAACGATCTCAAGTTTATTT | GGGGACCACTTTGTACAAGAAAGCTGGGTGCGGTAGCGTTTCTCATTAGTTTCTA |
| NCR343-3UTR-MsGW | GGGGACAGCTTTCTTGTACAAAGTGGGGATACAAAGAACGACGGAAAAAAATCA | GGGGACAACTTTGTATAATAAAGTTGGTGTGTTTCGTATCATCTCAAATCAT |
| pNCR-new35-MsGW | GGGGACAACTTTGTATAGAAAAGTTGGGTGTTGTAGAAGGAAGAGGAGGAG | GGGGACTGCTTTTTTGTACAAACTTGGGTAAAACAATTTTTTTAATCTCTTTATAAGTAAAC |
| NCR-new35-coding-MsGW | GGGGACAAGTTTGTACAAAAAAGCAGGCTGGATGCAAAGGAAGAAAAATATGGC | GGGGACCACTTTGTACAAGAAAGCTGGGTGAATAATAAACATTTCAGGTTGTCA |
| NCR-new35-3UTR-MsGW | GGGGACAGCTTTCTTGTACAAAGTGGGGATAATAAGTTACTTTAATAAGTAA | GGGGACAACTTTGTATAATAAAGTTGGCCGTTTTGACAGCTCTAACAGTA |
| pNCR211-MsGW | GGGGACAACTTTGTATAGAAAAGTTGGGCGGAGTGTAGGGGTGATGTTGTT | GGGGACTGCTTTTTTGTACAAACTTGGTCTTTTTTAACTTTGTATATAAC |
| NCR211-coding-MsGW | GGGGACAAGTTTGTACAAAAAAGCAGGCTGGATGGCTGAAATTCTCAAGTTTGT | GGGGACCACTTTGTACAAGAAAGCTGGGTGTAATTTCCATGAGAAGTAACGTG |
| NCR211-3UTR-MsGW | GGGGACAGCTTTCTTGTACAAAGTGGGGAATGTTATAAGTTGATCAAAGTC | GGGGACAACTTTGTATAATAAAGTTGGGAATACTGTGTTGTTATATAATACG |
| **MsGW clones in pKm43GW-rolD::EGFP vector for modification of cysteine residues of NCR peptides :** | | |
| NCR343-coding mC34S-MsGW | GGATGGATTAAATCTAAAGTCGATGAAGAT | ATCTTCATCGACTTTAGATTTAATCCATCC |
| NCR-new35-coding mC41S-MsGW | CAGCTTACATTCCTTCCATTTCTGATGATGA | TCATCATCAGAAATGGAAGGAATGTAAGCTG |
| NCR211-coding mC28S-MsGW | ACATTACAGGAGAGTCTGATACTGAC | GTCAGTATCAGACTCTCCTGTAATGT |
| **MsGW clones in pK7m34GW-rolD::EGFP vector for *in situ* localisation of NCR peptides :** | | |
| NCR343-coding-noSTOP-MsGW | GGGGACAAGTTTGTACAAAAAAGCAGGCTGGATGGCTAACGATCTCAAGTTTATTT | GGGGACCACTTTGTACAAGAAAGCTGGGTGCATGTATGGCTTTTTGGGTATCTCT |
| NCR-new35-coding-noSTOP-MsGW | GGGGACAAGTTTGTACAAAAAAGCAGGCTGGATGCAAAGGAAGAAAAATATGGC | GGGGACCACTTTGTACAAGAAAGCTGGGTGTCTTTGAGGTAAATCATCGTCC |
| **GW clones in pKGWFS7-prUBQ10::dsREd vector to monitor the activity of *NCR* promoters:** | | |
| pNCR343-GUS-GW | GGGGACAAGTTTGTACAAAAAAGCAGGCTGGCAACATTTTTCTCTCTTATGCCTTT | GGGGACCACTTTGTACAAGAAAGCTGGGTATTTGTTTTCTTTGTTTTTGTAACAAATAGG |
| pNCR-new35-GUS-GW | GGGGACAAGTTTGTACAAAAAAGCAGGCTTGTTGTAGAAGGAAGAGGAGGAG | GGGGACCACTTTGTACAAGAAAGCTGGGTGTAAAACAATTTTTTTAATCTCTTTATAAGTAAAC |
| pNCR-169-GUS-GW | GGGGACAAGTTTGTACAAAAAAGCAGGCTAAAACCGCTACAATCAGATCG | GGGGACCACTTTGTACAAGAAAGCTGGGTTGTTTTTCCCCTTTGCGTGAAA |
| pNCR-211-GUS-GW | GGGGACAAGTTTGTACAAAAAAGCAGGCTCGGAGTGTAGGGGTGATGTTGTT | GGGGACCACTTTGTACAAGAAAGCTGGGTGTCTTTTTTAACTTTGTATATAAC |
